# Evidence of selection in *UCP1* gene region suggests local adaptation to irradiance rather than cold temperatures in savanna monkeys (*Chlorocebus* spp.)

**DOI:** 10.1101/2022.06.28.496889

**Authors:** Christian M. Gagnon, Hannes Svardal, Anna J. Jasinska, Jennifer Danzy Cramer, Nelson B. Freimer, J. Paul Grobler, Trudy R. Turner, Christopher A. Schmitt

## Abstract

The genus *Chlorocebus* is widely distributed throughout sub-Saharan Africa, and in the last 300 thousand years expanded from equatorial Africa into the southernmost latitudes of the continent. In these new environments, colder climate was a likely driver of natural selection. We investigated population-level genetic variation in the mitochondrial uncoupling protein 1 (*UCP1*) gene region – implicated in non-shivering thermogenesis within brown/beige adipocytes – in 73 wild savanna monkeys from three taxa representing this southern expansion (*C. pygerythrus hilgerti, C. cynosuros, C. p. pygerythrus*) ranging from Kenya to South Africa. We found 17 SNPs with extended haplotype homozygosity consistent with positive selective sweeps, 10 of which show no significant LD with each other. Phylogenetic generalized least squares modeling with ecological covariates suggest that most derived allele frequencies are significantly associated with solar irradiance and winter precipitation, rather than overall low temperatures. This selection and association with irradiance appears to be driven by a population isolate in the southern coastal belt of South Africa. We suggest that sunbathing behaviors common to savanna monkeys, in combination with strength of solar irradiance, may mediate adaptations to thermal stress via non-shivering thermogenesis among savanna monkeys. The variants we discovered all lie in non-coding regions, some with previously documented regulatory functions, calling for further validation and research.

## Background

Cold environments pose a physiological challenge for homeothermic endotherms, which must find ways to conserve or produce heat, to maintain bodily functions. Severe or long-term exposure to cold temperatures can result in hypothermia or death, but there are also less obvious costs to warm-blooded species inhabiting cold environments. Even while surviving, energy allocated for successfully maintaining body temperature may diminish the energy available for other physiological needs, like reproduction (Bronson, 1985). Long term cold exposure may also slow the rate of biochemical reactions such as the production of hormones and enzymes (D’Amico et al., 2002), potentially indirectly affecting fitness, for example by mediating aspects of reproduction and pregnancy, growth and development, even behavior. Among endotherms, and mammals in particular, extreme cold has been an evolutionary problem that has resulted in several adaptive solutions, ranging from hibernation and torpor (Hochachka and Guppy 1987; Davenport, 1992), to shifts in body size and proportions meant to retain body heat (e.g., Bergmann’s and Allen’s Rules, Meiri & Dayan, 2003; Tilkens et al., 2007). However, there also exist shorter-term physiological responses that generate heat to offset losses in body temperature, such as shivering (Hohtola, 2004), shifts in circulation (Solonin and Katsyuba, 2003), and non-shivering thermogenesis.

Non-shivering thermogenesis (NST) is a short-term, heat generating phenotype mediated by brown adipose tissue (BAT). Brown adipocytes are metabolically active fat cells, the metabolism of which are driven by the expression of the *UCP1* gene (which codes for the protein UCP1, or thermogenin) (Cannon & Nedergaard, 2004). Thermogenin, a membrane protein, disrupts the electron transport chain at the inner mitochondrial membrane, effectively inhibiting adenosine triphosphate (ATP) synthesis and resulting in the production of heat (Cannon & Nedergaard, 2004). More specifically, UCP1 proteins embedded in the mitochondrial membrane interact with free fatty-acids (FA) in the intermembrane space, resulting in an increase in the ability of UCP1 to transport hydrogen ions across the mitochondrial membrane. The presence of FA in the intermembrane space disrupts the activity of ATP synthase directing more ions to UCP1 proteins in order to reach the mitochondrial matrix. Heat is produced as hydrogen ions pass through the uncoupling protein into the matrix, primarily as a byproduct of the metabolic breakdown of FAs. Although other genes may play a role in regulating NST in muscle tissue (Nowack et al., 2017; Nowack et al., 2019), *UCP1* has been shown to be critical for NST in brown adipose tissue, highlighting its adaptive significance (Golozoubova et al., 2006).

Cold-induced heat production via *UCP1-*mediated NST is believed to have played a critical role in the evolution and expansion of eutherian mammals (Hughes et al., 2009). The phylogenetic history of the mammalian *UCP1* gene shows that variants increasing NST function were evolutionarily favored among small-bodied mammals, and in larger mammals these variants may protect neonates, but NST is far from a universal solution to offsetting the cold, and several mammalian lineages have thrived while having lost the trait (Gaudry et al., 2018). Among primates - most notably in humans - NST appears to have been a critically important adaptation to climatic variation. Adaptations to cold climates, including NST, likely contributed to the successful migration out of Africa by early humans (Sazini et al., 2014) and the eventual colonization of more extreme cold environments in northern Eurasia (van Marken Lichtenbelt et al., 2009). Among humans, genotype variants of the *UCP1* gene have a significant impact on the degree of NST in the brown adipose tissue of healthy subjects (Nishimura et al., 2017). Although the evolution of *UCP1* thermoregulatory function in mammals and beyond has been the focus of previous studies (D’Amico et al., 2002; Mozo et al., 2005; Klingenspor et al., 2008; Hughes et al., 2009; Nishimura et al., 2017), we have a limited understanding of the selective pressures that have shaped this gene in the primate order.

Savanna monkeys (*Chlorocebus* spp.) are an excellent wild primate model in which to study thermoregulatory adaptations. The genus likely originated in equatorial central Africa and, with the exception of the rainforest dwelling dryas monkey (*Chlorocebus dryas*; van der Valk et al., 2020) and Bale monkey (*Chlorocebus djamdjamensis*; Mekonnen et al., 2018), are largely restricted to savanna environments. Genomic evidence suggests that, in the past ∼200-400 ky, savanna monkeys expanded from eastern equatorial Africa into the southern hemisphere (Warren et al., 2015; Svardal et al., 2017; van der Valk et al., 2020). Vervet monkeys (*Chlorocebus pygerythrus*, sensu lato; Turner et al., 2019) represent this southern expansion, with extant populations ranging from Ethiopia through South Africa along the Indian Ocean coast. South African vervets (*Chlorocebus pygerythrus pygerythrus*), in particular, have presented an interesting case study for cold adaptation as they regularly experience subzero temperatures at the southern extremes of their range (Danzy et al., 2012; McFarland et al., 2014). Within this range, vervets also occupy habitats at a variety of elevations, ranging from sea level to altitudes over 2,000 masl (Turner et al., 2018), with the higher elevations further exacerbating cold temperatures. Like our own species, vervet monkeys have successfully balanced adaptations to cold with the thermoregulatory demands of living in open savannahs, which can also reach very high temperatures (Lubbe et al., 2014). Furthermore, their relatively close phylogenetic relationship with *Homo sapiens* and wide geographic range make vervets a compelling model system for studying the evolutionary significance of NST in humans and primates more broadly.

To test this idea, we investigated variation in the *UCP1* gene region for signs of selection in three closely related taxa of savanna monkey, at various times all subsumed into the species *Chlorocebus pygerythrus*, representing what we refer to as the “southern expansion” of the genus into temperate latitudes. We chose vervet monkeys as our study species specifically because of their ability to inhabit environments that are uncharacteristically cold for most non-human primates, as well as their wide latitudinal, altitudinal, and climatic range. We hypothesized that as vervet monkeys expanded their geographic range into parts of southern Africa, exposure to colder temperatures would have favored *UCP1* variants which promote NST. Given their wide geographic distribution, we predicted generally strong differentiation within the gene region between populations sampled from areas geographically distant from one another, consistent with isolation by distance. However, we also predicted evidence of positive selection in restricted, functionally relevant regions of the *UCP1* gene region — including, potentially, in cis-acting regulatory regions — of populations sampled from higher latitudes more generally, and in geographic regions where climatic conditions are coldest, either generally (due to high elevations) or during particularly cold seasons in more temperate latitudes.

## Methods

### Study Populations

Data for this project were generated by the International Vervet Research Consortium from a sample of 163 savanna monkeys (*Chlorocebus* spp.) from 6 closely related taxa, captured as part of a larger sampling effort in 11 countries across Africa and the Caribbean, with particularly extensive sampling in South Africa (Jasinska et al., 2013; Svardal et al., 2017; Turner et al., 2019). We focused on 73 individuals from 3 vervet taxa: *Chlorocebus pygerythrus hilgerti, Chlorocebus cynosuros*, and *Chlorocebus pygerythrus pygerythrus* (Figure 1A; Supplemental Table 1). In this paper we adhere roughly to taxonomic distinctions made by Groves (2001), although recent nuclear and mitochondrial genomic evidence both suggest a relatively deep division suggesting a species-level distinction between *C. p. hilgerti* and *C. p. pygerythrus* (Svardal et al., 2017; Dolotovskaya et al., 2017), while evidence of gene flow between these two taxa and *C. cynosuros* suggest that the latter is nested within the larger *C. pygerythrus* clade (as first proposed by Dandelot, 1959). A taxonomic revision may be in order for the genus (Svardal et al., 2017), but is not within the scope of this paper. Each taxon was further subdivided into distinct populations, when possible, based on inferences from the whole-genome phylogeny (see below).

**Figure 1.**
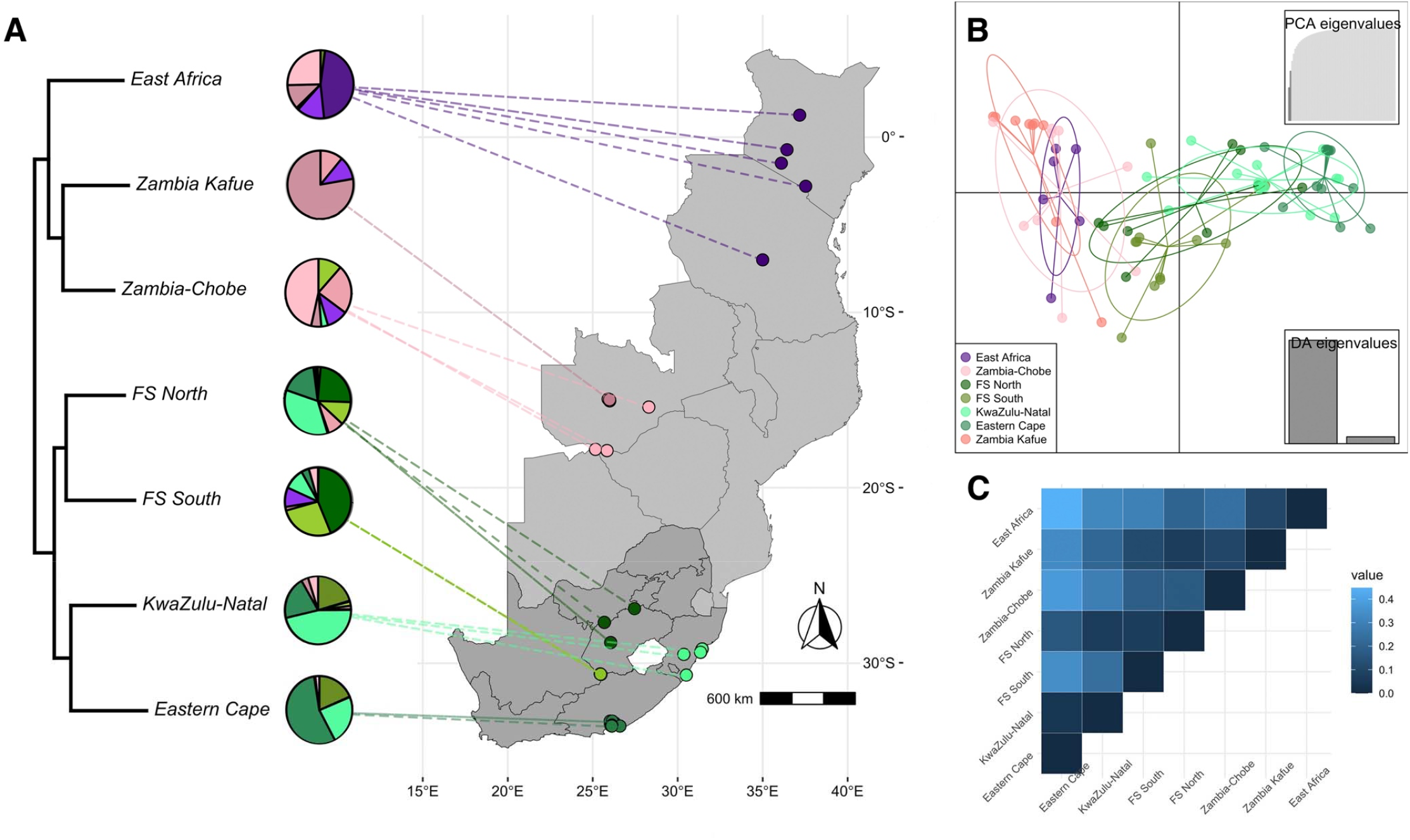
Population differentiation in the southern expansion of savanna monkeys in the *UCP1* gene region, including A) population relationships based on the whole-genome phylogeny of savanna monkeys, with pie charts indicating substructuring in the *UCP1* gene region in which purple clusters are primarily found in *C. p. hilgerti*, pink in *C. cynosuros* (darker pink = Zambia-Kafue; lighter pink = Zambia-Chobe), and shades of green in *C. p. pygerythrus* (dark green = Free State North; olive green = Free State South; light sea green = KwaZulu-Natal; dark sea green = Eastern Cape), with the map showing collection locations from southern and eastern Africa (South Africa, the more darkly shaded region, denotes regional boundaries, all others are national boundaries); B) DAPC results for the *UCP1* gene region; and C) F_ST_ heatmap for the *UCP1* gene region comparing defined populations in the southern expansion.

### Sequence Data

Whole genome sequences were previously generated by collaborators at the McDonnell Genome Institute at Washington University in St. Louis (Warren et al., 2015; Svardal et al., 2017), and analyzed in this study using a publicly available variant call format (VCF) file generated at the Gregor Mendel Institute (Svardal et al., 2017). We used *tabix* (Li, 2011) in *HTSlib v1*.*10*.*2* and *vcftools* (Danecek et al., 2011) to isolate a ∼28 kb gene region around *UCP1* (*ChlSab1*.*1* positions 7:87,492,195-87,502,665), including 10 kb upstream and downstream of the coding region itself to capture potential cis-regulatory regions.

### Whole-Genome Phylogeny

We used *SNPRelate v. 1*.*24*.*0* (Zheng et al., 2012) to generate FASTA alignments from the VCF file, and *phangorn* (*v2*.*5*.*5*; Schliep, 2011), *phytools* (*v0*.*6-99*; Revell, 2012), and *geiger* (*v2*.*0*.*6*.*4*; Pennell et al., 2014) to construct a neighbor-joining tree with a Jukes-Cantor mutation model representing whole-genome phylogenetic relationships for all 163 individuals in our original sample. We pruned this phylogeny to drop tips not represented in the study population sample.

### Population Structure

We used discriminant analyses of principal components (DAPCs) in *adegenet v*.*2*.*1*.*2* (Jombart, 2008; Jombart & Ahmed, 2011) and calculated the Fixation Index (F_ST_) using *hierfstat v*.*0*.*04-22* (Goudet, 2005) to model population structure. To statistically validate the patterns noted, we ran analyses of molecular variance (AMOVA) in *poppr v. 2*.*8*.*6* (Kanvar et al., 2014). In both DAPC and F_ST_ analyses we used sample population as a grouping variable, while in AMOVA we used population nested within taxon. In the F_ST_ analysis we used the *Nei87* setting, based on Nei’s (1987) method, to assess genetic distance between populations. We also ran principal component and admixture analysis using *LEA v. 3*.*0*.*0* (Frichot & François, 2015), using the *snmf* function with standard settings to visualize entropy criterion values to define *K* levels of population differentiation. To model isolation by distance, we assessed the correlation between genetic and geographic distance matrices in our sample using Mantel tests implemented in *vegan v. 2*.*5-6*, using the Pearson method (Oksanen et al., 2019). We constructed our genetic distance matrix in *poppr*, also using Nei’s method (Nei, 1978), and our geographic distance matrix from GPS points associated with trapping locations (Supplemental Table 1) using *raster v. 3*.*4-5* (Hijmans & van Etten, 2012).

### Assessing Selection

We calculated Hardy Weinberg Equilibrium (HWE) values with *pegas* (*v. 0*.*12*; Paradis, 2010) to identify candidate loci potentially experiencing selective forces, both within the whole southern expansion and in local populations. We assessed linkage disequilibrium (LD) with *gpart v*.*1*.*6*.*0* (Kim and Yoo, 2020), using standard *BigLD* setting and the *r*^*2*^ method (to avoid the “ceiling effect” common to using D’ in small samples; Marroni et al., 2011). SNPs considered to be in LD with each other (*r*^*2*^ > 0.85) were subsumed into a single representative locus for downstream analyses. We calculated site-frequency spectrum statistics for the whole gene region including Tajima’s D, and Fu and Li’s D* and F* (*PopGenome v*.*2*.*7*.*5*; Pfeifer et al., 2014, *pegas v*.*0*.*12*; Paradis, 2010), as well as a sliding window Tajima’s D in *vcftools* with a window size of 500 bp, for both the southern expansion and each constituent subpopulation in South Africa. We compared *UCP1* regional values of Tajima’s D and Fu and Li’s D* and F* to those from a sample of 1000 random, non-overlapping regions of equivalent size from vervet chromosome 7 to assess relative significance.

To calculate integrated haplotype scores (iHS) and assess extended haplotype homozygosity (EHH) both across the whole gene region and for selected *UCP1* loci, respectively, we inferred the ancestral allele sequence for our sample population using the program *Est-sfs v*.*2*.*03* using Kimura’s mutation model (Keightley & Jackson, 2018). We chose the rhesus macaque reference (*Macaca mulatta*; BCM Mmul_8.0.1/rheMac8) as our outgroup, which we downloaded using *biomaRt v. 2*.*44*.*4* (Durnick et al., 2009; Durnick et al., 2005). We aligned it to the the vervet reference genome (*Chlorocebus sabaeus*; Chlorocebus_sabeus 1.1/chlSab2) using *rMSA v. 0*.*99*.*0* (Hahsler & Manguy, 2020) and *MAFFT v. 7*.*467* (Katoh & Standley, 2013) using standard settings, and trimmed the macaque reference to the vervet extent visually in *JalView v. 2*.*11*.*1*.*3* (Waterhouse et al., 2009). Ancestral alleles were assigned to the major vervet allele when the probability assigned by *Est-sfs* was above 0.70, and to the minor allele if below 0.30. When the probability was between these benchmarks, we assigned the ancestral allele to the allele shared by the two outgroups if it matched the population allele; when there was no concordance between the two outgroups (n = 7), we chose the *Chlorocebus* reference allele. We used *rehh v*.*3*.*0*.*1* (Gautier et al., 2017) to estimate iHS and EHH using standard settings, but with a frequency bin of 0.15 to calculate iHS. We used a significance threshold of 2 for absolute iHS values (Voight et al., 2006) with a window size of 3000 bp and an overlap of 300 bp. We used a significance threshold value of 1.3 for considering individual SNPs for inclusion in further analyses.

### Ecological Covariates

We used GPS coordinates recorded at each trapping location to download altitude, annual mean temperature, mean temperature of the coldest month, winter precipitation levels, and mean temperature of the wet season, among other ecological covariates, for each population for the 10-year period from 2005-2010 from the *WorldClim2* online database (Fick & Hijmans, 2017). We also collected data on mean annual solar irradiance (measured in MJ/m^2^/day) for these points from the 10-year period from 2005-2010, originally generated by the NASA Langley Research Center (LaRC) POWER Project funded through the NASA Earth Science/Applied Science Program, using *nasapower 3*.*0*.*1* (Sparks, 2018). We standardized all covariates using z-scores, and then used the package *PerfomanceAnalytics v. 2*.*0*.*4* (Peterson et al., 2020) to reduce strongly correlated covariates. Correlations of standardized covariates, and the distributions of the final ecological covariate chosen for each population can be seen in Supplemental Figure 1.

### Modeling Allele Frequency by Ecological Covariates

We used phylogenetic generalized least squares (PGLS) regression, implemented in the package *nlme v. 3*.*1-150* (Pinheiro et al., 2021), to model variation in derived allele frequency of each target locus by geoclimatic variables including latitude, elevation, insolation/irradiation, annual mean temperature, mean temperature of the coldest month, and mean winter temperature. We incorporated our phylogenomic tree into our models using a Brownian correlation structure to account for average genetic distance across populations. We then used an information theoretic approach (Burnham & Anderson, 2002) to assess ecological covariate inclusion for each locus putatively experiencing selection, using the lowest Akaike Information Criterion modified for small sample sizes (AICc) to select the appropriate model.

### Data Availability

Raw sequence data are publicly available through the NCBI Sequence Read Archive (SRA) under BioProject numbers PRJNA168521, PRJNA168472, PRJNA168520, PRJNA168527 and PRJNA168522. The VCF file used in this study is available from the European Variation Archive (EVA) under accession PRJEB22988. The data used in this analysis and the associated analytical pipeline are publicly available via the Dryad data repository (Schmitt et al., 2022), including instructions for downloading and processing all relevant online datasets and code.

## Results

### Whole-Genome Phylogeny and Isolation by Distance

The maximum likelihood phylogeny we constructed accords with previously published assessments of these taxa (Warren et al., 2015; Svardal et al., 2017; Supplemental Figure 2), with *C. p. hilgerti* and *C. cynosuros* clustering separately from *C. p. pygerythrus*, presumably reflecting the more recent and relatively larger amount of gene flow estimated to have occurred between these two taxa (Svardal et al., 2017). The whole-genome phylogeny further suggests that the monkeys sampled in South Africa are best represented by two geographically and genetically distinct populations, which can each be further divided into two clusters: the Free State (represented by Free State North and Free State South), and the southern coastal belt, represented by vervets from KwaZulu-Natal (KZN) and the Eastern Cape (Figure 1A). One individual sampled at a rehabilitation center in Limpopo (VSAJ2008) appears to have been transplanted from the Free State North population, and so was assigned to that population for subsequent analyses. Another rehabilitant (VSAI3005) did not fit easily into any of these populations, instead appearing basal to the KZN/Eastern Cape cluster, and so was excluded from further analyses. The two individuals sampled in Botswana (VBOA1003, VBOA1005) and taxonomically assigned to *C. p. pygerythrus* appear to be genomically nested within *C. cynosuros*, and so we assigned them to that taxon for these analyses. We also divided *C. cynosuros* into two populations based on geographic and genetic distance. One clade from Kafue National Park in Zambia (Zambia Kafue) is nested within a larger clade comprised of both the Zambian Lusaka samples and Botswana samples from Chobe; we included the latter groups with those sampled in Livingstone as a population (Zambia-Chobe) due to similarities in climate and to retain sufficient sample size for downstream modeling (Figure 1A; Supplemental Figure 2).

### Population Structure

DAPC indicates clear genetic differentiation in the *UCP1* gene region among our study populations. Again in concordance with previous work on the whole genome (Warren et al., 2015; Svardal et al., 2017), we see relatively high genetic similarity in the *UCP1* gene region between *C. cynosuros* and *C. p. hilgerti* (presumably due to more recent or extensive introgression), with *C. p. pygerythrus* showing clear divergence. *C. p. pygerythrus* showed the widest range of variation (Figure 1B), to the extent that the South African southern coastal belt populations (Eastern Cape and KZN) show no overlap with *C. p. hilgerti/cynosuros*. AMOVA results confirm this pattern, showing significant variation between taxa (13.8%, df = 2, p < 0.05) and also between populations within taxa (12.6%, df = 4, p < 0.01) in the *UCP1* gene region (Supplemental Table 2; Supplemental Figure 3), with *C. p. pygerythrus* appearing to drive the former result and the latter driven by the separation between the Free State and southern coastal belt populations within South Africa. F_ST_ results are identical to those from DAPC, showing relatively high F_ST_ between the southern coastal belt population and all others, with the exception of relatively high similarity between KZN and Free State North (F_ST_ = 0.064) (Figure 1C; Supplemental Table 3).

Entropy criterion values from the principal component and admixture analyses strongly suggest isolation by distance across the southern expansion, with a population differentiation of K = 10 clusters (Supplemental Figure 4). The gradual geographic pattern of genetic differentiation and generally low taxon- or population-specific substructure in genetic clusters also suggest isolation by distance, although there are a few patterns of note in keeping with the analyses above, including distinct clusters associated with the South African southern coastal belt that are shared with Free State North, and large overlap in shared clusters between *C. p. hilgerti* and *C. cynosuros* (Figure 1A; Supplemental Figure 5). Mantel tests of Nei’s genetic distance between individual vervets compared to geographic distances between sample collection points show a strong positive correlation between genetic and geographic distances, again suggesting isolation by distance (Mantel’s *r* = 0.44, p = 0.001).

### Linkage Disequilibrium

We identified 14 linkage blocks in the *UCP1* gene region (Figure 2; Supplemental Figure 6; Supplemental Table 4), including a large block (B-09) encompassing both known 5’ enhancer regions, the basal promoter region, the associated CpG island, the 5’ UTR, Exon 1, and most of Intron 1-2 of *UCP1*. We also found clear linkage between several loci of interest (see below), and used LD to reduce the number of loci assessed for selection to 10 unlinked loci of interest (Supplemental Figure 7).

### Assessing Selection

Site-frequency spectrum statistics indicate no significant deviations from neutrality in the *UCP1* gene region, as a whole, in the southern expansion. Tajima’s D was overall positive (D_Tajima_ = 0.70), although in *C. p. pygerythrus* alone it was more strongly if not significantly positive (D_Tajima_ = 1.23). The only local population showing even a slightly negative, albeit insignificant, Tajima’s D was the Eastern Cape (D_Tajima_ = -0.021). Assessments of each statistic for 1000 random, non-overlapping regions of vervet chromosome 7 suggest that Fu and Li’s D* and F* are both within the normal range of values across the chromosome. Tajima’s D, however, is markedly positive compared to the rest of the chromosome (p = 0.060), again suggesting an excess of polymorphisms in the *UCP1* region. A sliding window assessment of Tajima’s D in the southern expansion indicates several regions with a strong excess of polymorphisms, including a 1000 bp window (7:87497500-87498500, D_Tajima_ = 2.04/2.33) in the 3’ UTR, a 500 bp window (7:87503000-87503500, D_Tajima_ = 2.04) in Intron 2-3, and a 500 bp sequence in the 5’ upstream region (7:87508500-87509000, D_Tajima_ = 2.84) (Supplemental Table 6a). Given we are discussing more localized selection processes, we also assessed individual populations in South Africa and found the same marked excess of polymorphisms, particularly upstream of the enhancer regions in Free State North (7:87507000-87509500; Supplemental Table 6b), and including the promotor and enhancer regions in Free State South (7:87501500-87509500; Supplemental Table 6c). This was less marked in the southern coastal belt populations (Supplemental Table 6d-e), particularly in Eastern Cape where the entire gene region showed negative Tajima’s D (Supplemental Table 6d), although only one 500 bp segment in the 5’ upstream region neared the significance threshold (7:87511000-87511500, D_Tajima_ = -2.00).

Seventeen SNPs met our threshold for consideration (p_iHS_ > 1.3) based on integrated haplotype score (iHS) in the southern expansion (Supplemental Table 7), with the most compelling evidence for selection appearing in the upstream region of the gene (Supplemental Figure 8). Of these loci, several showed compelling evidence suggesting positive selective sweeps based on visual inspections of extended haplotype homozygosity (Figure 3; Supplemental Figure 9). Several of these SNPs also failed to meet the expectations of Hardy Weinberg Equilibrium in the larger southern expansion or among local populations (p < 0.05) (Table 1).

**Figure 2.**
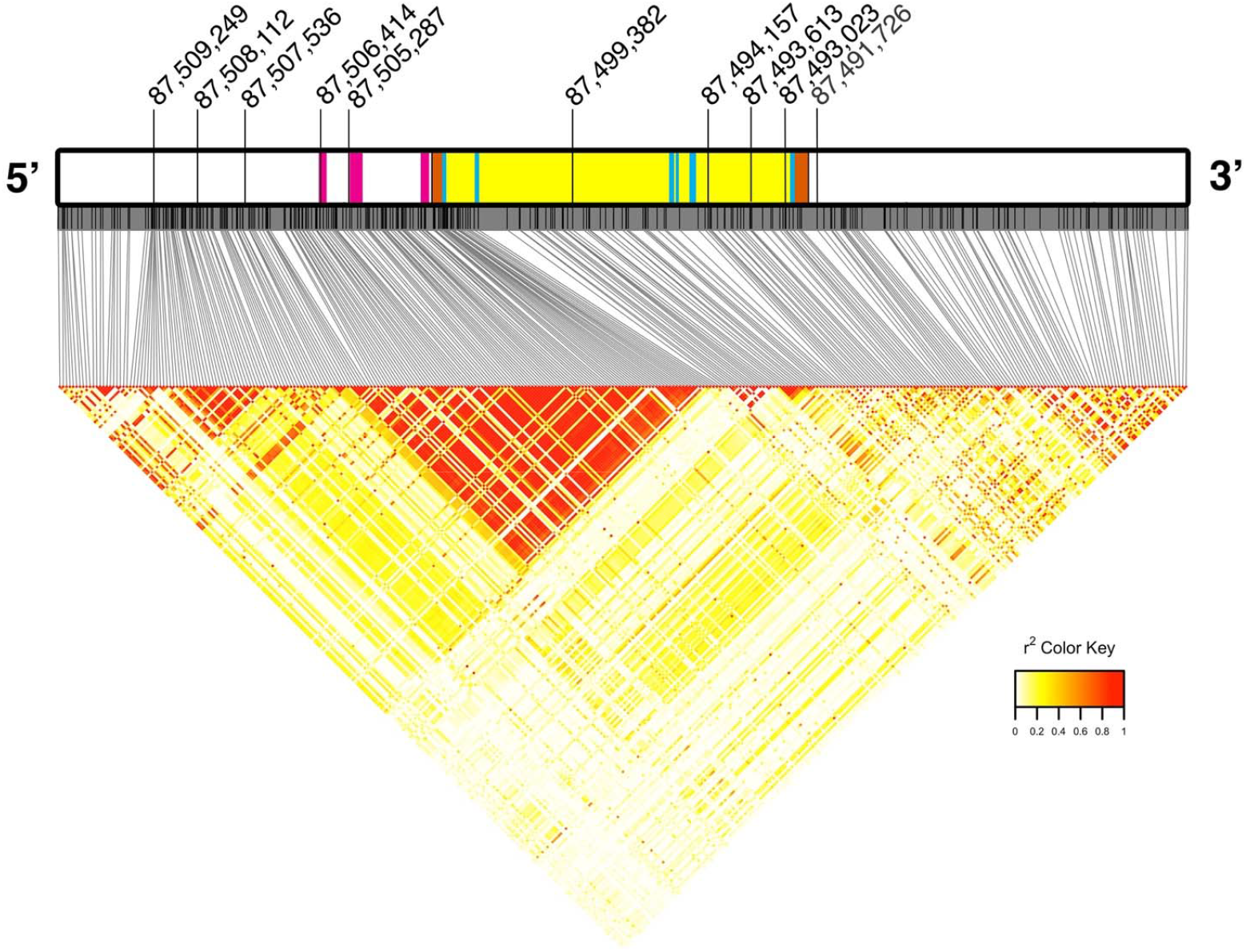
*UCP1* gene region highlighting candidate SNPs undergoing putative positive selective sweeps and LD heatblocks. Gene region includes *UCP1* (Positions 87,502,665 to 87,492,195, on the reverse strand) and 10 kilobases upstream and downstream. Thin black lines indicate SNPs located in the gene region, and those labeled above the gene diagram are SNPs undergoing putative positive selective sweeps (see Table 1 for details). In the gene diagram, magenta blocks represent 5’ upstream enhancers and the basal promoter region, orange blocks represent UTRs, blue blocks represent exons, and yellow blocks represent introns. The LD heatmap highlights the large LD block (B-09) encompassing the enhancers, promoter, 3’ UTR, and exon 1.

**Figure 3.**
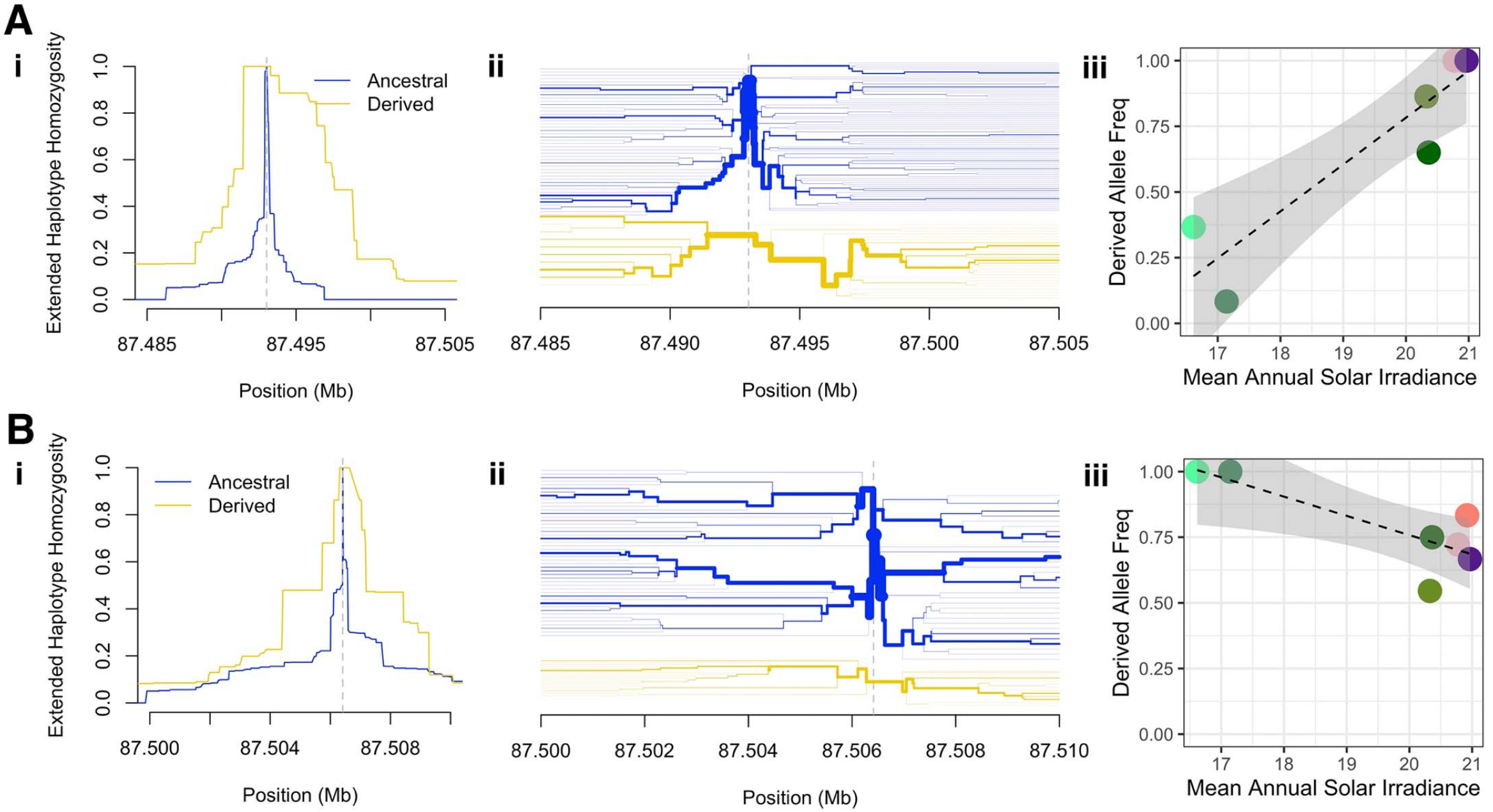
Examples of extended haplotype homozygosity (EHH) and ecological correlations with derived allele frequencies for A) 7:87493023 and b) 7:87506414 showing i) extended haplotype homozygosity around the candidate SNP, ii) bifurcation patterns around the candidate SNP, and iii) association of mean annual solar irradiation with derived allele frequencies among savanna monkey populations. In EHH diagrams, haplotypes around the ancestral allele are indicated in blue, and around the derived allele in yellow. For the plots of association between irradiance and population derived allele frequencies, population colors are the same as indicated in Figure 1. Similar figures for the remaining candidate SNPs are available in Supplemental Figures 9 and 10.

**Table 1.** Targeted SNPs in the *UCP1* gene region in the southern expansion and their associated indicators of selection. Tajima’s D values are from a 500 bp sliding window analysis conducted on either the South African (ZA) or southern expansion (SE) populations. Further details for each analysis/result are available in Supplemental Materials.

| SNP | Position | Polymorphism | Location | Block | Tajima's D |  | HWE |  | iHS | Best Model | Significant Ecological Covariates |  |  |  |  |  |
| --- | --- | --- | --- | --- | --- | --- | --- | --- | --- | --- | --- | --- | --- | --- | --- | --- |
| | | | | | ZA | SE | ZA | SE | | | Sig. Covariate(s) | $\beta$ | p-value | *r* | p-value | Outliers |
| 1 | 7:87491726 | *469C>T | 3' Downstream | B-02 | 0.57 | 0.78 | - | * | 1.28 | 1 + Winter Precip. + Irradiance | Winter Precipitation | 0.521 | 0.018 | 0.934 | 0.003 | EC/KZN |
|  |  |  |  |  |  |  |  |  |  |  | Irradiance | 0.607 | 0.011 |  |  |  |
| 2 | 7:87493023 | 6-304G>A | Intron 5-6 | B-02 | 1.79 | 1.00 | * | *** | 1.29 | 1 + Irradiance | Irradiance | 0.310 | 0.011 | 0.920 | 0.003 | EC/KZN |
| 3 | 7:87493613 | 6-895C>T | Intron 5-6 | B-03 | 0.45 | -0.51 | - | - | 1.34 | 1 + Winter Precip. | Winter Precipitation | -0.262 | 0.024 | 0.831 | 0.021 |  |
| 4 | 7:87494157 | 5+743C>T | Intron 5-6 | B-03 | 1.25 | -0.24 | - | - | 1.85 | Null | - | - | - | - | - |  |
| 5 | 7:87499382 | 2+2137T>C | Intron 2-3 | B-06 | -1.41 | -1.37 | - | . | 1.57 | Null | - | - | - | - | - |  |
| 6 | 7:87505287 | -2622C>A | 5' Enhancer | B-09 | 1.47 | 0.95 | - | - | 1.72 | 1 + Winter Precip. | Winter Precipitation | 0.162 | 0.073 | 0.796 | 0.032 |  |
| 7 | 7:87506414 | -3749A>T | 5' Enhancer | B-09 | 1.39 | 0.91 | - | - | 2.84 | 1 + Irradiance | Irradiance | -0.150 | 0.031 | 0.812 | 0.026 | EC/KZN |
| 8 | 7:87507536 | -4871T>C | 5' Upstream | B-10 | -0.07 | 1.06 | - | *** | 1.84 | Null | - | - | - | - | - |  |
| 9 | 7:87508112 | -5447C>G | 5' Upstream | B-11 | 1.40 | 1.56 | - | - | 2.00 | 1 + Min. Temp. Coldest Month | Min. Temp. Coldest Month | 0.125 | 0.000 | 0.731 | 0.062 |  |
| 10 | 7:87509248 | -6583C>A | 5' Upstream | B-14 | 2.66 | 1.02 | - | . | 2.72 | 1 + Irradiance | Irradiance | -0.094 | 0.000 | 0.982 | 0.000 | EC/KZN |

### Modeling Genotype by Ecological Covariates

We targeted ten SNPs for PGLS analysis based on iHS/EHH and cumulative evidence of selection (Table 1). Upon inspection of correlations between potential ecological covariates, we reduced them to include only normalized values for annual mean temperature (bio1), minimum temperature of the coldest month (bio6), mean temperature of the wet season (bio8), winter precipitation levels (bio19) and mean annual solar irradiance (insol). Using our model reduction approach, we found several environmental variables to be significantly associated with derived alleles at *UCP1* loci showing potential for having undergone recent selection, including mean annual solar irradiance, winter precipitation levels, and the minimum temperature of the coldest month (Table 1; Figure 3; Supplemental Figure 10).

## Discussion

We hypothesized that the *UCP1* gene would have undergone selection in relation to the expansion of *Chlorocebus* into more temperate geographic areas where temperatures regularly fall below freezing in the winter months, reflecting selection on NST as a way of offsetting these colder temperatures. This gene region generally shows differentiation consistent with both phylogenetic and geographic distance, and much of the variation therein is consistent with isolation by distance; however, there is clear divergence in excess of isolation by distance in South African *C. p. pygerythrus* populations when compared to the rest of the genus (Figure 1).

This divergence in South Africa is particularly marked in the coastal belt south and east of the Drakensberg mountains, represented in this study by populations sampled in KwaZulu-Natal and the Eastern Cape. These populations show high F_ST_ relative to all other populations, share relatively unique haplotypes to the exclusion of most other populations, and show similar patterns of variation relative to ecogeographic variables (Figure 1; Figure 3), while also showing the strongest evidence for positive selective sweeps (see below). Strong differentiation of these populations from the rest of the South African vervets has been noted previously in subspecies taxonomy (which differentiated them as *Chlorocebus pygerythrus cloeti*; Groves, 2001), studies of SIV_agm_ differentiation within South African vervets (Ma et al., 2013), and studies of mtDNA haplotypes (Turner et al., 2016). Turner et al. (2016) suggest that the Tugela River in the northeast and limited dispersal abilities across the Nama Karoo in the southwest have resulted in this coastal belt being a population isolate, potentially subject to strong drift. Our results here further suggest that unique patterns of selection could also be a factor in the differentiation of *UCP1* in these populations, given that the southern coastal belt is as cold as the rest of our savanna monkey sample while also showing relatively low levels of solar irradiance (due to being at comparatively low elevations). The extent to which migration is constrained between the southern coastal belt populations and the rest of *C. p. pygerythrus* in South Africa is important, particularly given theoretical work demonstrating that isolated populations with constrained migration are more likely to see locally beneficial alleles increase in frequency in response to ecological variation (e.g., Yeaman & Otto, 2011). The apparently high amount of shared variation between the FS North and KwaZulu-Natal populations in this gene region (Figure 1) suggest recent/frequent migration, and further calls for greater sampling in this area to suss out the relative contributions of drift and selection to these results.

Both haplotype- and site frequency-based analyses suggest that several SNPs in the *UCP1* region may have undergone recent selection among savanna monkeys. Many of the SNPs identified in this study appear to be in a large linkage block (B-09; Figure 2; Supplemental Figure 6; Supplemental Table 4) encompassing the *UCP1* basal promoter region and associated CpG island, suggesting that these variants may have undergone positive selection related to some regulatory effect influencing *UCP1* expression. Indeed, SNPs in this upstream region in humans have previously been associated with increased *UCP1* expression, resulting in greater body heat generation when exposed to cold (in particular the A-3826G variant, among others; e.g., Rose et al., 2011; Nishimura et al., 2017). Positive Tajima’s D results, primarily in the 5’ upstream region among local populations in the Free State, suggest either balancing selection or decreases in population size. The only populations where we observe the negative values of Tajima’s D associated with positive selective sweeps are in the southern coastal belt, and particularly in the Eastern Cape.

It is important to note that these statistics indicating selection can be influenced by demographic parameters that we are unable to account for in this study. For example, site frequency spectrum–based statistics like Tajima’s D and Fu & Li’s D* and F* can be strongly influenced by demographic patterns, with the genomic signature of population expansions, in particular, mirroring that of positive selective sweeps (Weigand & Leese, 2018). Although we generally did not find significant results using these statistics in the *UCP1* gene region (which is not unusual for recent selection; Sabeti et al., 2002), inferred recent increases in effective population sizes in *C. p. pygerythrus*, and smaller increases in *C. cynosuros*, could interfere with these results (Warren et al., 2015). To suss out the difference between positive selection and background selection using these site-frequency spectrum-based methods would require a better understanding of, for example, demographic history and recombination rate differences between these populations, which is not currently feasible (Stephan, 2010). For the linkage-based methods used here, strong bottlenecks and high migration between populations may reduce power to detect positive selective sweeps (Weigand & Leese, 2018). Even with the high admixture previously reported among these populations (Svardal et al., 2017) we were able to detect positive selective sweeps. These methods are also generally not optimal for detecting soft selective sweeps (Weigand & Leese, 2018), although it is difficult to infer how much standing ancestral variation in *UCP1* may have been available in savanna monkeys prior to the southern expansion given the general dearth of annotated whole genomes in closely related cercopithecine primates (Hernandez et al., 2020). Finally, linkage-based methods, in particular, perform poorly when populations are hierarchical in structure (Vatsiou et al., 2016; Weigand & Leese, 2018), which is a concern given the clear hierarchical structure and inferred localized migration and introgression patterns in this sample (Svardal et al., 2017).

PGLS models indicate that the derived allele frequencies of many of these variants are associated with geoclimatic variables that indicate selection in relation to cold. Some associations, like increases in derived allele frequencies with the minimum temperature of the coldest month (7:87508112) and increases (7:87491726, 7:87505287) or decreases (7:87493613) with winter precipitation, suggest that these *UCP1* variants may be under selection by modifying NST to offset the cold directly (with cold wet fur, as in winter rain conditions, being a particularly difficult thermoregulatory challenge; e.g., Cristóbal-Azkarate et al., 2016). A surprisingly common association was with the strength of mean annual solar irradiance, with derived allele frequencies strongly associated with *weaker* irradiance (i.e., 7:87506414, 7:87509248). Given that sunbathing has previously been identified as an important behavior for offsetting cold temperatures in vervet monkeys (Danzy et al., 2012; Figure 4), we propose that vervet populations exposed to relatively low levels of solar radiation - such as those in the low-elevation South African southern coastal belt - receive little thermoregulatory benefit from sunbathing behaviors, making adaptive changes in NST (via variation in *UCP1* function/expression) necessary as a means of increasing thermoregulatory resistance to the cold. Loci 7:87506414 (along with linked locus 7:87503019) and 7:87505287 are particularly compelling as they reside within or near upstream regulatory regions: 7:87503019 is less than 100 bp upstream of the basal promoter region (Shore et al., 2013), 7:87506414 is in the enhancer region identified by Gaudry & Campbell (2017) between the relatively conserved CRE-3 and RARE-1 motifs, while 7:87505287 lies 5 bp upstream of the conserved enhancer region identified by Shore et al. (2013). These three loci may be especially good candidates for functional validation for expression-based differences in *UCP1* phenotypes in savanna monkeys.

**Figure 4.**
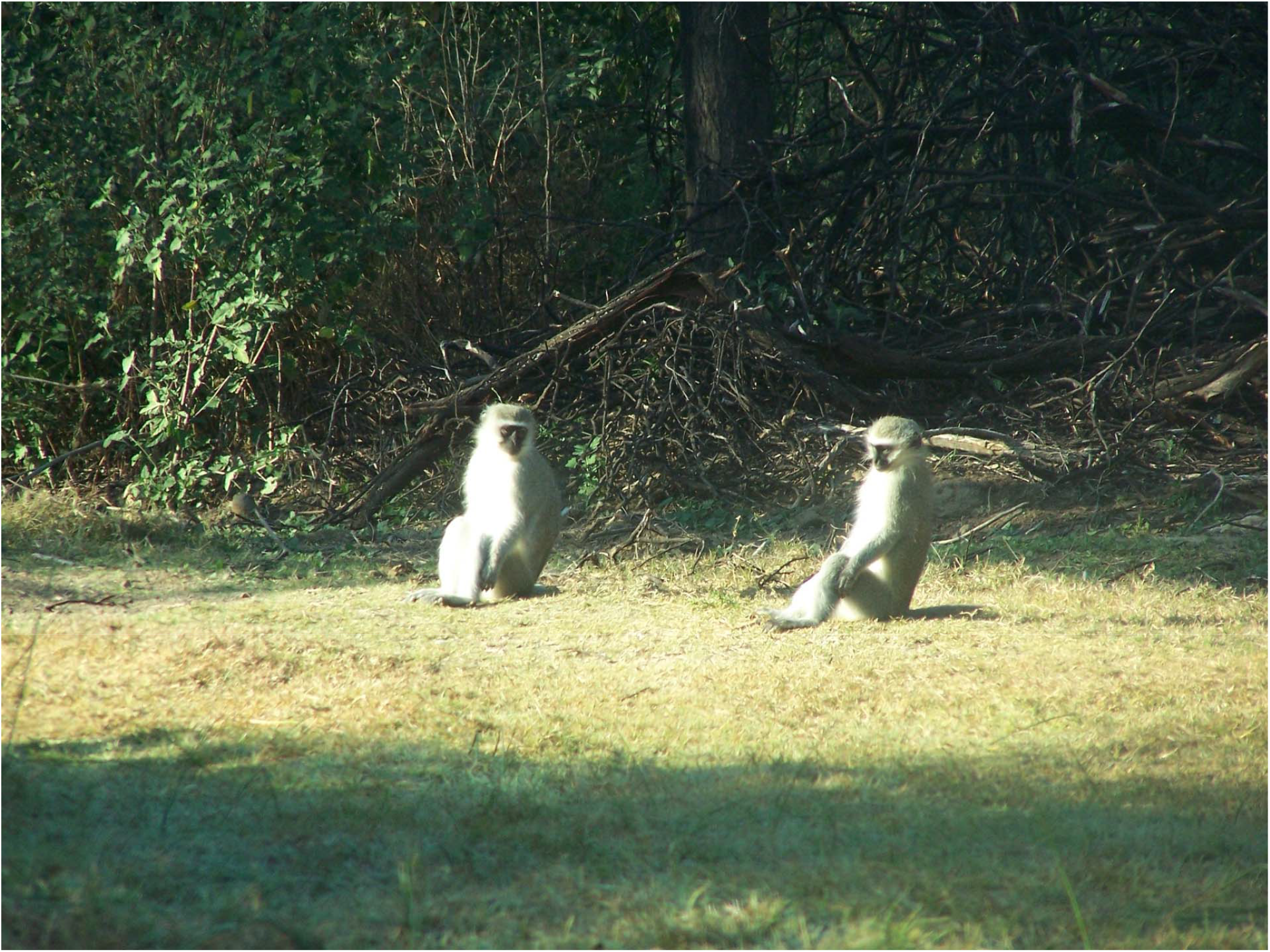
Sunbathing vervet monkeys (*Chlorocebus pygerythrus pygerythrus*) in Soetdoring Nature Reserve, Free State, South Africa. This photo was previously published by Danzy et al. (2012) and is reproduced with permission from *African Primates*.

In contrast to the negative association between mean annual solar irradiance and derived allele frequencies in the 5’ upstream region of *UCP1*, we note that loci 7:87491756 and 7:87493023 show strongly positive associations with irradiance, with these loci showing the southern coastal belt populations as outliers. In these cases, our hypothesized pattern for the derived allele appears to be inverted in that the signs of selection strongly point towards greater EHH and putative selection in what are likely warmer conditions. It may simply be the case that these alleles are simply linked with the actual allele of effect, making the direction of association arbitrary. In the case of 7:87491756, the evidence for EHH appears relatively weak (Supplemental Figure 9a), and the relationship between the derived allele and irradiance seems to be exclusively driven by the southern coastal belt (Supplemental Figure 10a); elimination of these outliers would eliminate the association with irradiance, but would also strengthen the relationship of derived allele frequencies with winter precipitation, creating a more significantly positive association suggesting selection in *C. p. hilgerti* populations. In the case of 7:87493023, the derived allele appears to be fixed in populations of *C. cynosuros* and *C. p. hilgerti*, where it shows strong EHH (Supplemental Figures 9b and 10b); given that it eliminates one of the only CpG sites in Intron 5-6, it’s possible that the selection is somehow related to methylation potential at this locus and its effects on *UCP1* expression. It is also possible this is an artifact of selection on other sequence variants in the larger linkage block (B-02) in this region of *UCP1*, which includes the functionally relevant 3’ UTR (Mayr, 2019). Aside from these potential functional explanations, it is unclear why this pattern is being observed.

Although the evidence of selection at certain loci in the *UCP1* gene region and the apparent relationship between their allele frequencies and geoclimatic variables are compelling, there are many ways in which this study could be strengthened to better inform this question. Our sample sizes should be adequately informative given past studies on wild non-model organisms (i.e., Nazareno et al., 2017; Satkoski Trask et al., 2011); however, increased sampling, particularly across geoclimatic variables and within each population (particularly *C. cynosuros* and *C. p. hilgerti*) would strengthen our confidence in these results. Additionally, although we did not include elevation as a covariate in our PGLS models (due to a 95% correlation with irradiance, which we considered to be more biologically meaningful for this study than elevation alone), work in mice has suggested that hypoxia may also independently influence NST by significantly reducing thermogenic capacity (Beaudry & McClellan., 2010). Given that the southern coastal belt populations are the only populations in our sample near sea-level, increased variation in elevation in our sample at different latitudes might be necessary to disentangle a potential relationship between elevation, irradiance, and NST. Given the archived data available from the International Vervet Research Consortium (Jasinska et al., 2013; Turner et al., 2019), this could be attainable with the resequencing of already-sampled individuals not included in this dataset. Further sampling in areas to increase geoclimatic diversity and in populations that bridge taxonomic and geographic gaps in our dataset - such as *C. p. pygerythrus* in Mozambique and Malawi, *C. p. rufoviridis* in Malawi and Tanzania, and *C. p. pygerythrus* in central and southern Botswana - could also provide greater information on the relative strength of selection and drift in the southern expansion.

To test explicitly in the wild whether behaviors like sunbathing may be implicated in selection for NST-related variants associated with irradiance would require an investigation of such behaviors in tandem with thermoregulatory physiology across relevant genotypes. Given previous research, it is likely that other behavioral factors beyond sunbathing also mediate the adaptive value of such variants (i.e., diet/feeding, dominance status and social integration, McFarland et al., 2014, 2015; sex, body size, residency status, social networks: Henzi et al., 2017; body position and microhabitat selection: Gestich et al., 2014; clustering and embracing behaviors: Gartlan & Brain, 1968). Incorporating genotype data at key loci into already-existing biophysical modeling systems of thermoregulation in wild savanna monkeys (i.e., Mathewson et al., 2020) could be a relatively simple and effective means of testing these relationships.

One important question raised by this study is whether the variants we identified actually have a functional effect on the expression of *UCP1* and NST. As has been shown in human and mouse models, upstream variants of *UCP1* may yield significant differences in the thermogenic potential of brown fat, making those loci found in upstream enhancer and promotor regions (i.e., 7:87503019, 7:87505287, 7:87506414) particularly compelling candidates for validation. However, we presently know very little regarding how the thermoregulatory function of brown adipose tissue is regulated in non-human primates. Quantifying changes in vervet *UCP1* expression associated with the variants identified here, either through adipose biopsy in captive individuals or through assessing relative *UCP1* expression in biobanked samples that have the variants in question (such as those collected by the IVRC; Jasinska et al., 2013; Turner et al., 2019), would help us to better characterize and study this critically important adaptive mammalian feature. Such studies of localized adaptive change in UCP1 expression may inform how our hominin ancestors managed to rapidly expand their geographical range beyond equatorial Africa, as well as how contemporary non-human primates may endure the thermoregulatory challenges of our changing global climate.

## Acknowledgements

The authors gratefully acknowledge the many people with whom they have worked over the years to obtain and process these data. Lisa Nevell provided the original spark for this idea while teaching about human *UCP1* at BU. Many students in the Sensory Morphology and Anthropological Genomics Lab at Boston University helped to refine this genomics pipeline at various times during its development, including especially Becca DeCamp, Gianna Grob and Erica Sun, as well as Mansa Asiedu, Ishrat Chowdhury, Tennyson Cooper, Jane Hilsenrath, Alexia Lancea, Kimberly Louisor, Diem Maxwell, Elizabeth Shelton, and Akshata Shukla (some with funding by the BU Undergraduate Research Opportunities Program). Original data collection included assistance in the field by Fred Brett in Ethiopia; Nicholas Dracopoli, James Else and The Institute for Primate Research in Kenya; Oliver “Pess” Morton and Yoon Jung at UCLA; numerous field assistants in South Africa including especially Riel Coetzer, Magali Jacquier, Helene DeNys, JD Pampush, Elzet Answegen, Tegan Gaetano, Dewald DuPlessis, Micah Beller, David Beller and Pess Morton, with support by the University of Limpopo and the University of the Free State. Sampling was conducted under permits and permissions issued by the following: the Botswana Ministry of Environment & Wildlife and Tourism the Ethiopian Wildlife Conservation Authority; Ministry of Tourism and Wildlife, Kenya; in South Africa the Department of Economic Development and Environmental Affairs, Eastern Cape; Department of Tourism, Environmental and Economic Affairs, Free State Province; Ezemvelo KZN Wildlife; Department of Economic Development, Environment and Tourism, Limpopo; the Department of Agriculture, Conservation and Environment, Mpumalanga; the Department of Environment and Nature Conservation, Northern Cape; the South African National Department of Environmental Affairs; and the Zambia Wildlife Authority. We thank J. Brenchley, K. Reimann (R24OD010976), and J. Baulu and the Barbados Primate Research Center and Wildlife Reserve for providing samples of Tanzanian origin. The authors also acknowledge their appreciation to all the veterinarians who worked with them to safely obtain samples, especially Lizanne Meiring and Murray Stokoe. This work was funded by Boston University, and the original collections work funded by The Fulbright Foundation, NSF (SOC 74-24166, BNS 770-3322, BCS 0938969) NIH (R01RR0163009), the University of Wisconsin-Milwaukee, The University of the Free State, the University of Limpopo, and the Coriell Institute.

## CRediT Author Statement

**Christian M. Gagnon**: methodology, formal analysis, software, writing - original draft and editing, visualization; **Hannes Svardal, Anna J. Jasinska**: resources, data curation, writing - editing. **Jennifer Danzy Cramer**: writing - editing, photos. **Nelson B. Freimer**: resources, funding acquisition. **J. Paul Grobler, Trudy R. Turner**: resources, funding acquisition, writing - editing. **Christopher A. Schmitt**: conceptualization, methodology, formal analysis, software, writing - original draft and editing, project administration, visualization.

## References

Beaudry, J. L. & McClelland, G. B. (2010). Thermogenesis in CD-1 mice after combined chronic hypoxia and cold acclimation. Comparative Biochemistry and Physiology Part B: Biochemistry and Molecular Biology, 157, 301–309.

Bronson, F. H. (1985). Mammalian reproduction: an ecological perspective. Biology of reproduction, 32(1), 1–26.

Cannon, B., & Nedergaard, J. A. N. (2004). Brown adipose tissue: function and physiological significance. Physiological reviews, 84(1), 277–359.

Cristóbal-Azkarate, J., Maréchal, L., Semple, S., Majolo, B., & MacLarnon, A. (2017). Metabolic strategies in wild male Barbary macaques: evidence from faecal measurement of thyroid hormone. Biology Letters, 12(4), 12(4), 20160168.

D’Amico, S., Claverie, P., Collins, T., Georlette, D., Gratia, E., Hoyoux, A., Meuwis, M-A., Feller, G., & Gerday, C. (2002). Molecular basis of cold adaptation. Philosophical Transactions of the Royal Society of London. Series B: Biological Sciences, 357(1423), 917–925.

Dandelot, P. (1959). Note sur la classification des Cercopitheques du groupe aethiops. Mammalia, 23, 357–368.

Danecek, P., Auton, A., Abecasis, G., Albers, C. A., Banks, E., DePristo, M. A., Handsaker, R. E., Lunter, G., Marth, G. T., Sherry, S. T., & McVean, G. (2011). The variant call format and VCFtools. Bioinformatics, 27(15), 2156–2158.

Danzy, J., Grobler, J. P., Freimer, N., & Turner, T. R. (2012). Sunbathing: a behavioral response to seasonal climatic change among South African vervet monkeys (Chlorocebus aethiops). African Primates, 7(2), 230–237.

Davenport, J. 1992. Animal Life at Low Temperature. Chapman & Hall, London.

Dolotovskaya, S., Torroba Bordallo, J., Haus, T., Noll, A., Hofreiter, M., Zinner, D., & Roos, C. (2017). Comparing mitogenomic timetrees for two African savannah primate genera (Chlorocebus and Papio). Zoological Journal of the Linnean Society, 181(2), 471–483.

Durinck, S., Spellman, P. T., Birney, E., & Huber, W. (2009). Mapping identifiers for the integration of genomic datasets with the R/Bioconductor package biomaRt. Nature protocols, 4(8), 1184.

Durinck, S., Moreau, Y., Kasprzyk, A., Davis, S., De Moor, B., Brazma, A., & Huber, W. (2005). BioMart and Bioconductor: a powerful link between biological databases and microarray data analysis. Bioinformatics, 21(16), 3439–3440.

Feldmann, H. M., Golozoubova, V., Cannon, B., & Nedergaard, J. (2009). UCP1 ablation induces obesity and abolishes diet-induced thermogenesis in mice exempt from thermal stress by living at thermoneutrality. Cell metabolism, 9(2), 203–209.

Fick, S. E., & Hijmans, R. J. (2017). WorldClim 2: new 1LJkm spatial resolution climate surfaces for global land areas. International journal of climatology, 37(12), 4302–4315.

Frichot, E., & François, O. (2015). LEA: An R package for landscape and ecological association studies. Methods in Ecology and Evolution, 6(8), 925–929.

Gartlan, J.S. & C.K. Brain. 1968. Ecology and social variability in Cercopithecus aethiops and C. mitis. In: Primates. Studies in Adaptation and Variability. P.C. Jay, ed. Rinehart and Winston, New York, NY. Pp. 253–292.

Gaudry, M. J. & Campbell, K. L. (2017). Evolution of UCP1 transcriptional regulatory elements across the mammalian phylogeny. Frontiers in Physiology, 8, 670.

Gaudry, M. J., Campbell, K. L., & Jastroch, M. (2018). Evolution of UCP1. Brown Adipose Tissue, 127–141.

Gautier, M., Klassmann, A., & Vitalis, R. (2017). rehh 2.0: a reimplementation of the R package rehh to detect positive selection from haplotype structure. Molecular ecology resources, 17(1), 78–90.

Gestich, C. C., Caselli, C. B., & Setz, E. Z. F. (2014). Behavioural thermoregulation in a small Neotropical primate. Ethology, 120(4), 331–339.

Golozoubova, V., Cannon, B., & Nedergaard, J. (2006). UCP1 is essential for adaptive adrenergic nonshivering thermogenesis. American Journal of Physiology-Endocrinology and Metabolism, 291(2), E350–E357.

Goudet, J. (2005). Hierfstat, a package for R to compute and test hierarchical FLJstatistics. Molecular Ecology Notes, 5(1), 184–186.

Groves, C.P. (2001). Primate Taxonomy. Smithsonian Institution Press, Washington DC.

Hahsler, M., & Manguy, J. (2020). rMSA: Interface for Popular Multiple Sequence Alignment Tools. R package version 0.99.0.

Henzi, S.P., Hetem, R., Fuller, A., Maloney, S., Young, C., Mitchell, D., Barrett, L., & McFarland, R. (2017). Consequences of sex-specific sociability for thermoregulation in male vervet monkeys during winter. Journal of zoology, 302(3), 193–200.

Hernandez, M. Shenk, M. K., & Perry, G. H. (2020). Factors influencing taxonomic unevenness in scientific research: a mixed-methods case study of non-human primate genomic sequence data generation. Royal Society Open Science, 7(9), 201206.

Hijmans, R. J., & van Etten, J. (2012). raster: Geographic analysis and modeling with raster data. R package version 2.0-12.

Hochachka, P. W., & Guppy, M. (1987). Metabolic arrest and the control of biological time. Harvard University Press.

Hohtola, E. (2004). Shivering thermogenesis in birds and mammals. In Life in the cold: evolution, mechanisms, adaptation, and application. 12th International Hibernation Symposium (pp. 241–252). Institute of Arctic Biology.

Hughes, D. A., Jastroch, M., Stoneking, M., & Klingenspor, M. (2009). Molecular evolution of UCP1 and the evolutionary history of mammalian non-shivering thermogenesis. BMC evolutionary biology, 9(1), 4.

Jasinska, A.J., Schmitt, C.A., Service, S.K., Cantor, R.M., Dewar, K., Jentsch, J.D., Kaplan, J.R., Turner, T.R., Warren, W.C., Weinstock, G.M. and Woods, R.P. (2013). Systems biology of the vervet monkey. ILAR journal, 54(2), 122–143.

Jombart, T. (2008). adegenet: a R package for the multivariate analysis of genetic markers. Bioinformatics, 24(11), 1403–1405.

Jombart, T., & Ahmed, I. (2011). adegenet 1.3-1: new tools for the analysis of genome-wide SNP data. Bioinformatics, 27(21), 3070–3071.

Kamvar, Z. N., Tabima, J. F., & Grünwald, N. J. (2014). Poppr: an R package for genetic analysis of populations with clonal, partially clonal, and/or sexual reproduction. PeerJ, 2, e281.

Katoh, K., & Standley, D. M. (2013). MAFFT multiple sequence alignment software version 7: improvements in performance and usability. Molecular biology and evolution, 30(4), 772–780.

Keightley, P. D., & Jackson, B. C. (2018). Inferring the probability of the derived vs. the ancestral allelic state at a polymorphic site. Genetics, 209(3), 897–906.

Klingenspor, M., Fromme, T., Hughes, D.A., Manzke, L., Polymeropoulos, E., Riemann, T., Trzcionka, M., Hirschberg, V. and Jastroch, M. (2008). An ancient look at UCP1. Biochimica et Biophysica Acta (BBA)-Bioenergetics, 1777(7-8), 637–641.

Li, H. (2011). Tabix: fast retrieval of sequence features from generic TAB-delimited files. Bioinformatics, 27(5), 718–719.

Lubbe, A., Hetem, R. S., McFarland, R., Barrett, L., Henzi, P. S., Mitchell, D., Meyer, L. C. R., · Maloney, S. K. & Fuller, A. (2014). Thermoregulatory plasticity in free-ranging vervet monkeys, Chlorocebus pygerythrus. Journal of Comparative Physiology B, 184(6), 799–809.

Ma, D., Jasinska, A., Kristoff, J., Grobler, P., Turner, T., Jung, Y., Schmitt, C., Raehtz, K., Martinez, N., Wijewardana, V., Tracy, R., Pandrea, I., Freimer, N., & Apetrei, C. (2013). SIVagm infection in wild African green monkeys from South Africa: epidemiology, natural history, and evolutionary considerations. PLoS Pathogens, 9(1), 1–18.

Marroni, F., Pinosio, S., Zaina, G., Fogolari, F., Felice, N., Cattonaro, F., & Morgante, M. (2011). Nucleotide diversity and linkage disequilibrium in Populus nigra cinnamyl alcohol dehydrogenase (CAD4) gene. Tree genetics & genomes, 7, 1011–1023.

Mathewson, P., Porter, W.P., Barrett, L., Fuller, A., Henzi, S.P., Hetem, R.S., Young, C., McFarland, R. (2020). Field data confirm the ability of a biophysical model to predict wild primate body temperature. Journal of thermal biology, 94, 102754.

Mayr, C. (2019). What are 3’ UTRs doing? Cold Spring Harbor Perspectives in Biology,11(10), a034728.

McFarland, R., Barrett, L., Boner, R., Freeman, N. J., & Henzi, S. P. (2014). Behavioral flexibility of vervet monkeys in response to climatic and social variability. American journal of physical anthropology, 154(3), 357–364.

McFarland, R., Fuller, A., Hetem, R.S., Mitchell, D., Maloney, S.K., Henzi S.P., & Barrett, L. (2015). Social integration confers thermal benefits in a gregarious primate. Journal of animal ecology, 84(3), 871–878.

Meiri, S., & Dayan, T. (2003). On the validity of Bergmann’s rule. Journal of biogeography, 30(3), 331–351.

Mekonnen, A., Rueness, E.K., Stenseth, N.C., Fashing, P.J., Bekele, A., Hernandez-Aguilar, R.A., Missbach, R., Haus, T., Zinner, D. and Roos, C. (2018). Population genetic structure and evolutionary history of Bale monkeys (Chlorocebus djamdjamensis) in the southern Ethiopian Highlands. BMC evolutionary biology, 18(1), 1–15.

Mozo, J., Emre, Y., Bouillaud, F., Ricquier, D., & Criscuolo, F. (2005). Thermoregulation: what role for UCPs in mammals and birds?. Bioscience reports, 25(3-4), 227–249.

Nazareno, A.G., Bemmels, J.B., Dick, C.W., and Lohmann, L.G. (2017). Minimum sample sizes for population genomics: an empirical study from an Amazonian plant species. Molecular ecology resources, 17(6), 1136–1147.

Nishimura, T., Katsumura, T., Motoi, M., Oota, H., & Watanuki, S. (2017). Experimental evidence reveals the UCP1 genotype changes the oxygen consumption attributed to non-shivering thermogenesis in humans. Scientific reports, 7(1), 5570.

Nei, M. (1978). Estimation of average heterozygosity and genetic distance from a small number of individuals. Genetics, 89(3), 583–590.

Nei, M. (1987). Molecular evolutionary genetics. Columbia university press.

Nowack, J., Giroud, S., Arnold, W., & Ruf, T. (2017). Muscle non-shivering thermogenesis and its role in the evolution of endothermy. Frontiers in physiology, 8, 889.

Nowack, J., Vetter, S.G., Stalder, G., Painer, J., Kral, M., Smith, S., Le, M.H., Jurcevic, P., Bieber, C., Arnold, W. and Ruf, T. (2019). Muscle nonshivering thermogenesis in a feral mammal. Scientific reports, 9(1), 1–10.

Obenchain, V., Lawrence, M., Carey, V., Gogarten, S., Shannon, P., & Morgan, M. (2014). VariantAnnotation: a Bioconductor package for exploration and annotation of genetic variants. Bioinformatics, 30(14), 2076–2078.

Oksanen, J., Blanchet, F.G., Kindt, R., Legendre, P., Minchin, P.R., O’hara, R.B., Simpson, G.L., Solymos, P., Stevens, M.H.H. and Wagner, H. (2019). Community ecology package. R package version, 2.5-6.

Paradis, E. (2010). pegas: an R package for population genetics with an integrated–modular approach. Bioinformatics, 26(3), 419–420.

Pennell, M.W., Eastman, J.M., Slater, G.J., Brown, J.W., Uyeda, J.C., FitzJohn, R.G., Alfaro, M.E. and Harmon, L.J. (2014). geiger v2. 0: an expanded suite of methods for fitting macroevolutionary models to phylogenetic trees. Bioinformatics, 30(15), 2216–2218.

Pfeifer, B., Wittelsbürger, U., Ramos-Onsins, S. E., & Lercher, M. J. (2014). PopGenome: an efficient Swiss army knife for population genomic analyses in R. Molecular biology and evolution, 31(7), 1929–1936.

Pinheiro, J., Bates, D., DebRoy, S., Sarkar, D., R Core Team (2021). nlme: Linear and Nonlinear Mixed Effects Models. R package version 3.1-152.

Revell, L. J. (2012). phytools: Phylogenetic tools for comparative biology (and other things) Methods Ecol. Evol, 3, 217–223.

Rose, G., Crocco, P., D’Aquila, P., Montesanto, A., Bellizi, D., and Passarino, G. (2011). Two variants in the upstream enhancer region of human UCP1 gene affect gene expression and are correlated with human longevity. Experimental gerontology, 46(11), 897–904.

Rothwell, N. J., & Stock, M. J. (1979). A role for brown adipose tissue in diet-induced thermogenesis. Nature, 281(5726), 31–35.

Sabeti, P., Reich, D., Higgins, J., Levine, H. Z. P., Richter, D. J., Schaffner, S. F., Gabriel, S. B., Platko, J. V., Patterson, N. J., McDonald, G. J., Ackerman, H. C., Campbell, S. J., Altschuler, D., Cooper, R., Kwiatkowski, D., Ward, R., & Lander, E. S. (2002). Detecting recent positive selection in the human genome from haplotype structure. Nature, 419, 832–837.

Satkoski Trask, J.A., Malhi, R., Kanthaswamy, S., Johnson, J., Garnica, W.T., Malladi, V.S., and Glenn Smith, D. (2011). The effect of SNP discovery method and sample size on estimation of population genetic data for Chinese and Indian rhesus macaques (Macaca mulatta). Primates, 52(2), 129–138.

Sazzini, M., Schiavo, G., De Fanti, S., Martelli, P. L., Casadio, R., & Luiselli, D. (2014). Searching for signatures of cold adaptations in modern and archaic humans: hints from the brown adipose tissue genes. Heredity, 113(3), 259–267.

Schliep, K. P. (2011). phangorn: phylogenetic analysis in R. Bioinformatics, 27(4), 592–593.

Schmitt, C. A., Gagnon, C. M., Svardal, H., Jasinska, A. J., Danzy Cramer, J., Freimer, N. B., Grobler, J. P., and Turner, T. R. (2022). Savanna monkey (Chlorocebus spp.) population genetics/genomics pipeline. Dryad, Dataset. doi: 20.5062/dryad.k3j9kd59z

Shore, A., Karamitri, A., Kemp, P., Speakman, J. R., and Lomax, M. A. (2010). Role of UCP1 enhancer methylation and chromatin remodeling in the control of UCP1 expression in murine adipose tissue. Diabetologia, 51, 1164–1173.

Shore, A., Emes, R. D., Wessely, F., Kemp, P., Cillo, C., D’Armiento, M., Hoggard, N., & Lomax, M. A. (2013). A comparative approach to understanding tissue-specific expression of uncoupling protein 1 expression in adipose tissue. Frontiers in Genetics, 3:304. doi: 10.3389/fgene.2012.00304

Solonin, Y. G., & Katsyuba, E. A. (2003). Thermoregulation and blood circulation in adults during short-term exposure to extreme temperatures. Human Physiology, 29(2), 188–194.

Sparks, A. H. (2018). nasapower: a NASA POWER global meteorology, surface solar energy and climatology data client for R.

Stephan, W. (2010). Genetic hitchhiking versus background selection: the controversy and its implications. Philosophical Transactions of the Royal Society B: Biological Sciences, 365(1544), 1245–1253.

Svardal, H., Jasinska, A.J., Apetrei, C., Coppola, G., Huang, Y., Schmitt, C.A., Jacquelin, B., Ramensky, V., Müller-Trutwin, M., Antonio, M. and Weinstock, G. (2017). Ancient hybridization and strong adaptation to viruses across African vervet monkey populations. Nature genetics, 49(12), 1705–1713.

Svardal, H., Jasinska, A. J., Schmitt, C. A., Huang, Y., Weinstock, G., Grobler, J. P., Wilson, R. K., Warren, W. C., Freimer, N. B., Nordberg, M., & Turner, T. R. (2018). Population genomics disentangles taxonomic relationships and identifies ancient hybridization in the genus Chlorocebus. American Journal of Physical Anthropology, 162(S64), 50.

Tilkens, M. J., Wall-Scheffler, C., Weaver, T. D., & Steudel-Numbers, K. (2007). The effects of body proportions on thermoregulation: an experimental assessment of Allen’s rule. Journal of human evolution, 53(3), 286–291.

Trayhurn, P. (1979). Thermoregulation in the diabetic-obese (db/db) mouse. The role of non-shivering thermogenesis in energy balance. Pflugers Archiv: European journal of physiology, 380(3), 227–232.

Turner, T.R., Coetzer, W.G., Schmitt, C.A., Lorenz, J.G., Freimer, N.B., & Grobler, J.P. (2016). Localized population divergence of vervet monkeys (Chlorocebus spp.) in South Africa: Evidence from mtDNA. American Journal of Physical Anthropology 159, 17–30.

Turner, T.R., Schmitt, C.A., Cramer, J.D., Lorenz, J., Grobler, J.P., Jolly, C.J., & Freimer, N.B. (2018). Morphological variation in the genus Chlorocebus: Ecogeographic and anthropogenically mediated variation in body mass, postcranial morphology, and growth. American journal of physical anthropology, 166, 682–707.

Turner, T.R., Schmitt, C.A., Cramer, J.D. (2019). Savanna Monkeys: The Genus Chlorocebus. Cambridge: Cambridge University Press.

van der Valk, T., Gonda, C.M., Silegowa, H., Almanza, S., Sifuentes-Romero, I., Hart, T.B., Hart, J.A., Detwiler, K.M. & Guschanski, K. (2020). The genome of the endangered dryas monkey provides new insights into the evolutionary history of the vervets. Molecular biology and evolution, 37(1), 183–194.

van Marken Lichtenbelt, W.D., Vanhommerig, J.W., Smulders, N.M., Drossaerts, J.M., Kemerink, G.J., Bouvy, N.D., Schrauwen, P. & Teule, G.J. (2009). Cold-activated brown adipose tissue in healthy men. New England Journal of Medicine, 360(15), 1500–1508.

Vatsiou, A.I., Bazin, E., & Gaggiotti, O.E. (2016). Detection of selective sweeps in structured populations: a comparison of recent methods. Molecular ecology, 25, 89–103.

Voight, B. F., Kuduravalli, S., Wen, X., & Pritchard, J. K. (2006). A map of recent positive selection in the human genome. PLOS Biology, 4(3), e72.

Warren, W.C., Jasinska, A.J., García-Pérez, R., Svardal, H., Tomlinson, C., Rocchi, M., Archidiacono, N., Capozzi, O., Minx, P., Montague, M.J. & Kyung, K. (2015). The genome of the vervet (Chlorocebus aethiops sabaeus). Genome research, 25(12), 1921–1933.

Waterhouse, A. M., Procter, J. B., Martin, D. M., Clamp, M., & Barton, G. J. (2009). Jalview Version 2— a multiple sequence alignment editor and analysis workbench. Bioinformatics, 25(9), 1189–1191.

Weigand, H., & Leese, F. (2018). Detecting signatures of positive selection in non-model species using genomic data. Zoological journal of the Linnean Society, 184(2), 528–583.

Yeaman, S., Otto, S.P. (2011). Establishment and maintenance of adaptive genetic divergence under migration, selection, and drift. Evolution, 67(7), 2123–2129.

Zheng, X., Levine, D., Shen, J., Gogarten, S., Laurie, C., & Weir, B. (2012). “A High-performance Computing Toolset for Relatedness and Principal Component Analysis of SNP Data.” Bioinformatics, 28(24), 3326–3328. doi: 10.1093/bioinformatics/bts606.

