## Supplemental Materials for "Evidence of selection in *UCP1* gene region suggests local adaptation to irradiance rather than cold temperatures in savanna monkeys (*Chlorocebus* spp.)"

**Supplementary Materials**

**Supplemental Table 1**: Genomic sample representing the southern expansion of savanna monkeys from equatorial to southern Africa, taken from the International Vervet Research Consortium dataset first published in Svardal et al. (2017). Latitude and longitude are in decimal degrees. South African locations taken by GPS, all others estimated from trapping location descriptions expect for Tanzanian samples, which are assigned a central location due to having no information regarding provenience.

| **Sample ID** | **Taxon** | **Country** | **Population** | **Location** | **Latitude** | **Longitude** |
| --- | --- | --- | --- | --- | --- | --- |
| AG23 | *hilgerti* | Tanzania | East Africa | Unknown | -7 | 35 |
| AG5417 | *hilgerti* | Tanzania | East Africa | Unknown | -7 | 35 |
| VBOA1003 | *cynosuros* | Botswana | Zambia-Chobe | Kasane | -17.81667 | 25.16 |
| VBOA1005 | *cynosuros* | Botswana | Zambia-Chobe | Kasane | -17.81667 | 25.16 |
| VKA3 | *hilgerti* | Kenya | East Africa | Samburu | 1.2571 | 37.1768 |
| VKB7 | *hilgerti* | Kenya | East Africa | Mosiro | -1.4833 | 36.1 |
| VKC6 | *hilgerti* | Kenya | East Africa | Naivasha | -0.7172 | 36.431 |
| VKD7 | *hilgerti* | Kenya | East Africa | Kimana | -2.8 | 37.5333 |
| VSAA2010 | *pygerythrus* | South Africa | FS North | Soetdoring | -28.821944 | 26.0594444 |
| VSAA2015 | *pygerythrus* | South Africa | FS North | Soetdoring | -28.821944 | 26.0594444 |
| VSAA2020 | *pygerythrus* | South Africa | FS North | Soetdoring | -28.821944 | 26.0594444 |
| VSAB1003 | *pygerythrus* | South Africa | FS South | Gariep | -30.634167 | 25.4869444 |
| VSAB2009 | *pygerythrus* | South Africa | FS South | Gariep | -30.606667 | 25.4463889 |
| VSAB2010 | *pygerythrus* | South Africa | FS South | Gariep | -30.606667 | 25.4463889 |
| VSAB2011 | *pygerythrus* | South Africa | FS South | Gariep | -30.606667 | 25.4463889 |
| VSAB2012 | *pygerythrus* | South Africa | FS South | Gariep | -30.606667 | 25.4463889 |
| VSAB2017 | *pygerythrus* | South Africa | FS South | Gariep | -30.606667 | 25.4463889 |
| VSAB2023 | *pygerythrus* | South Africa | FS South | Gariep | -30.606667 | 25.4463889 |
| VSAB3001 | *pygerythrus* | South Africa | FS South | Gariep | -30.606667 | 25.4463889 |
| VSAB3004 | *pygerythrus* | South Africa | FS South | Gariep | -30.606667 | 25.4463889 |
| VSAB5004 | *pygerythrus* | South Africa | FS South | Gariep | -30.616667 | 25.4622222 |
| VSAB5005 | *pygerythrus* | South Africa | FS South | Gariep | -30.616667 | 25.4622222 |
| VSAC1004 | *pygerythrus* | South Africa | FS North | Sandveld | -27.676111 | 25.6816667 |
| VSAC1012 | *pygerythrus* | South Africa | FS North | Sandveld | -27.676111 | 25.6816667 |
| VSAC1014 | *pygerythrus* | South Africa | FS North | Sandveld | -27.676111 | 25.6816667 |
| VSAC1015 | *pygerythrus* | South Africa | FS North | Sandveld | -27.676111 | 25.6816667 |
| VSAC1016 | *pygerythrus* | South Africa | FS North | Sandveld | -27.676111 | 25.6816667 |
| VSAD1003 | *pygerythrus* | South Africa | FS North | Parys | -26.893861 | 27.458 |
| VSAE2005 | *pygerythrus* | South Africa | KwaZulu-Natal | Zinkwazi | -29.207778 | 31.4241667 |
| VSAE2009 | *pygerythrus* | South Africa | KwaZulu-Natal | Zinkwazi | -29.207778 | 31.4241667 |
| VSAE2011 | *pygerythrus* | South Africa | KwaZulu-Natal | Zinkwazi | -29.207778 | 31.4241667 |
| VSAE3001 | *pygerythrus* | South Africa | KwaZulu-Natal | Zinkwazi | -29.185833 | 31.4416667 |
| VSAE3002 | *pygerythrus* | South Africa | KwaZulu-Natal | Zinkwazi | -29.185833 | 31.4416667 |
| VSAE3003 | *pygerythrus* | South Africa | KwaZulu-Natal | Zinkwazi | -29.185833 | 31.4416667 |
| VSAF1004 | *pygerythrus* | South Africa | KwaZulu-Natal | Blythedale | -29.374722 | 31.3488889 |
| VSAF1009 | *pygerythrus* | South Africa | KwaZulu-Natal | Blythedale | -29.374722 | 31.3488889 |
| VSAF1011 | *pygerythrus* | South Africa | KwaZulu-Natal | Blythedale | -29.374722 | 31.3488889 |
| VSAF1012 | *pygerythrus* | South Africa | KwaZulu-Natal | Blythedale | -29.374722 | 31.3488889 |
| VSAF1015 | *pygerythrus* | South Africa | KwaZulu-Natal | Blythedale | -29.374722 | 31.3488889 |
| VSAG2001 | *pygerythrus* | South Africa | KwaZulu-Natal | Kwela | -29.493536 | 30.360675 |
| VSAG2003 | *pygerythrus* | South Africa | KwaZulu-Natal | Kwela | -29.493536 | 30.360675 |
| VSAG2005 | *pygerythrus* | South Africa | KwaZulu-Natal | Kwela | -29.493536 | 30.360675 |
| VSAH1001 | *pygerythrus* | South Africa | KwaZulu-Natal | Anerley | -30.669722 | 30.5077778 |
| VSAI3005 | *pygerythrus* | South Africa | NA | Letsitele | NA | NA |
| VSAJ2008 | *pygerythrus* | South Africa | FS North | Letsitele | NA | NA |
| VSAK3004 | *pygerythrus* | South Africa | Eastern Cape | Shamwari | -33.473611 | 26.0422222 |
| VSAL1001 | *pygerythrus* | South Africa | Eastern Cape | Shamwari | -33.408889 | 26.1011111 |
| VSAL2002 | *pygerythrus* | South Africa | Eastern Cape | Shamwari | -33.288056 | 26.0313889 |
| VSAL3005 | *pygerythrus* | South Africa | Eastern Cape | Shamwari | -33.3975 | 26.2933333 |
| VSAL4002 | *pygerythrus* | South Africa | Eastern Cape | Shamwari | -33.293611 | 26.1483333 |
| VSAL5001 | *pygerythrus* | South Africa | Eastern Cape | Shamwari | -33.293611 | 26.1483333 |
| VSAM0021 | *pygerythrus* | South Africa | Eastern Cape | Amakhala | -33.5708 | 26.6025 |
| VSAM1003 | *pygerythrus* | South Africa | Eastern Cape | Amakhala | -33.536389 | 26.0694444 |
| VSAM2001 | *pygerythrus* | South Africa | Eastern Cape | Amakhala | -33.5025 | 26.1311111 |
| VSAM3001 | *pygerythrus* | South Africa | Eastern Cape | Amakhala | -33.550278 | 26.1247222 |
| VSAM4001 | *pygerythrus* | South Africa | Eastern Cape | Amakhala | -33.572222 | 26.1238889 |
| VSAM5007 | *pygerythrus* | South Africa | Eastern Cape | Addo | NA | NA |
| VZA1001 | *cynosuros* | Zambia | Zambia Kafue | Gate | -15.047083 | 25.9998667 |
| VZA1002 | *cynosuros* | Zambia | Zambia Kafue | Gate | -15.047083 | 25.9998667 |
| VZA1003 | *cynosuros* | Zambia | Zambia Kafue | Gate | -15.047083 | 25.9998667 |
| VZA1004 | *cynosuros* | Zambia | Zambia Kafue | Gate | -15.047083 | 25.9998667 |
| VZA2005 | *cynosuros* | Zambia | Zambia Kafue | Gate | -14.942733 | 25.9060167 |
| VZA2006 | *cynosuros* | Zambia | Zambia Kafue | Gate | -14.942733 | 25.9060167 |
| VZA3008 | *cynosuros* | Zambia | Zambia Kafue | Mukambi | -14.978317 | 25.9932167 |
| VZA3009 | *cynosuros* | Zambia | Zambia Kafue | Mukambi | -14.978317 | 25.9932167 |
| VZA3010 | *cynosuros* | Zambia | Zambia Kafue | Mukambi | -14.978317 | 25.9932167 |
| VZA4012 | *cynosuros* | Zambia | Zambia-Chobe | Lilayi | -15.4 | 28.3 |
| VZA4013 | *cynosuros* | Zambia | Zambia-Chobe | Lilayi | -15.4 | 28.3 |
| VZC1014 | *cynosuros* | Zambia | Zambia-Chobe | Livingstone | -17.884633 | 25.83925 |
| VZC1015 | *cynosuros* | Zambia | Zambia-Chobe | Livingstone | -17.884633 | 25.83925 |
| VZC1017 | *cynosuros* | Zambia | Zambia-Chobe | Livingstone | -17.884633 | 25.83925 |
| VZC1018 | *cynosuros* | Zambia | Zambia-Chobe | Livingstone | -17.884633 | 25.83925 |
| VZC1020 | *cynosuros* | Zambia | Zambia-Chobe | Livingstone | -17.884633 | 25.83925 |

**Supplemental Figure 1**: Final standardized ecological covariate a) distributions for each savanna monkey population in the southern expansion sample, and b) correlations among variables to determine those included in final models.

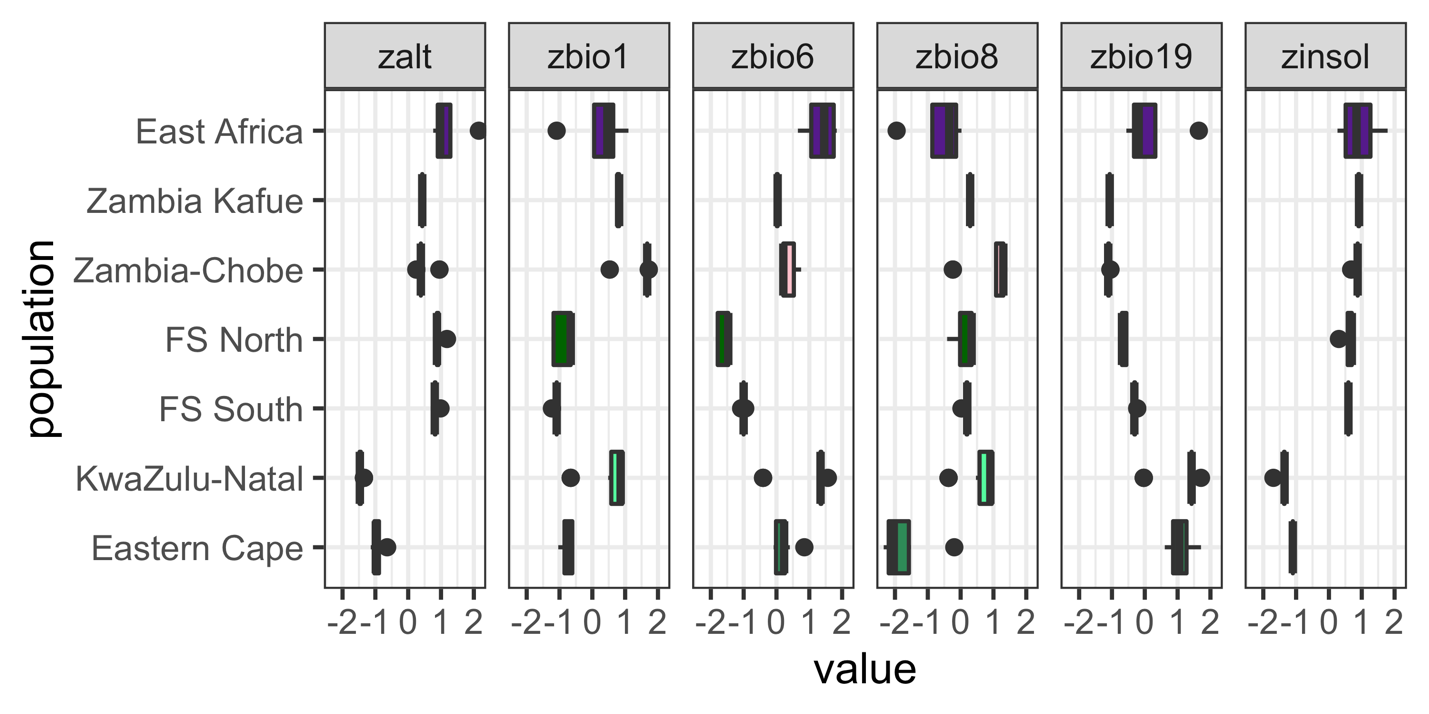

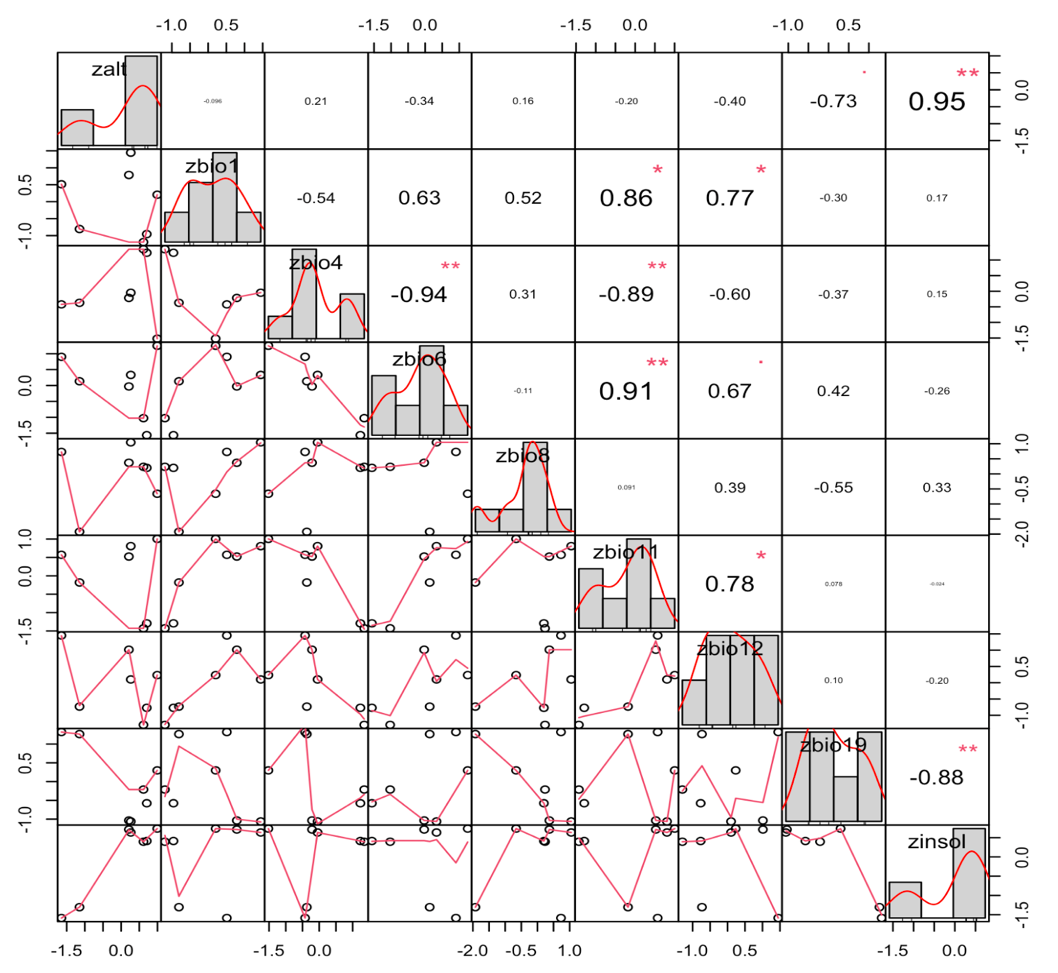

**Supplemental Figure 2**: Full whole genome phylogeny of the southern expansion sample. Tree uses the same coloring convention for populations as in Figure 1. Samples compressed for clarity include those from *C. tantalus* (red), *C. aethiops* (red), and *C. sabaeus* from The Gambia (orange) and St. Kitts & Nevis (yellow). All samples are described fully Svardal et al. (2017).

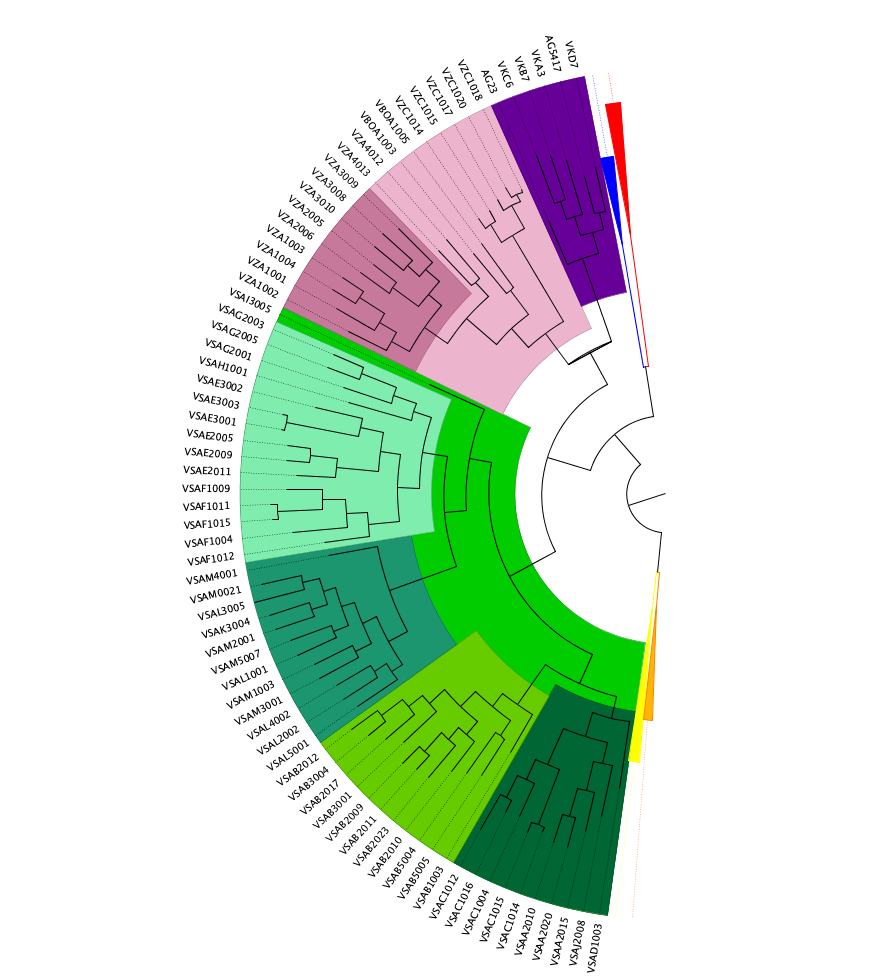

**Supplemental Table 2**: AMOVA results from {poppr}, showing significant differences between savanna monkey populations and taxa in the *UCP1* gene region.

| **Test** | **Obs.** | **Std. Obs.** | **Alter. Hyp.** | **p** |
| --- | --- | --- | --- | --- |
| Variation within samples | 46.42 | -3.70 | less | 0.01 |
| Variation between samples | -0.52 | -0.32 | greater | 0.66 |
| Variation between populations | 7.87 | 7.73 | greater | 0.01 |
| Variation between taxa | 8.61 | 2.22 | greater | 0.05 |

**Supplemental Figure 3**: Visualization of AMOVA results from {poppr}, showing significant differences between savanna monkey populations and taxa in the *UCP1* gene region.

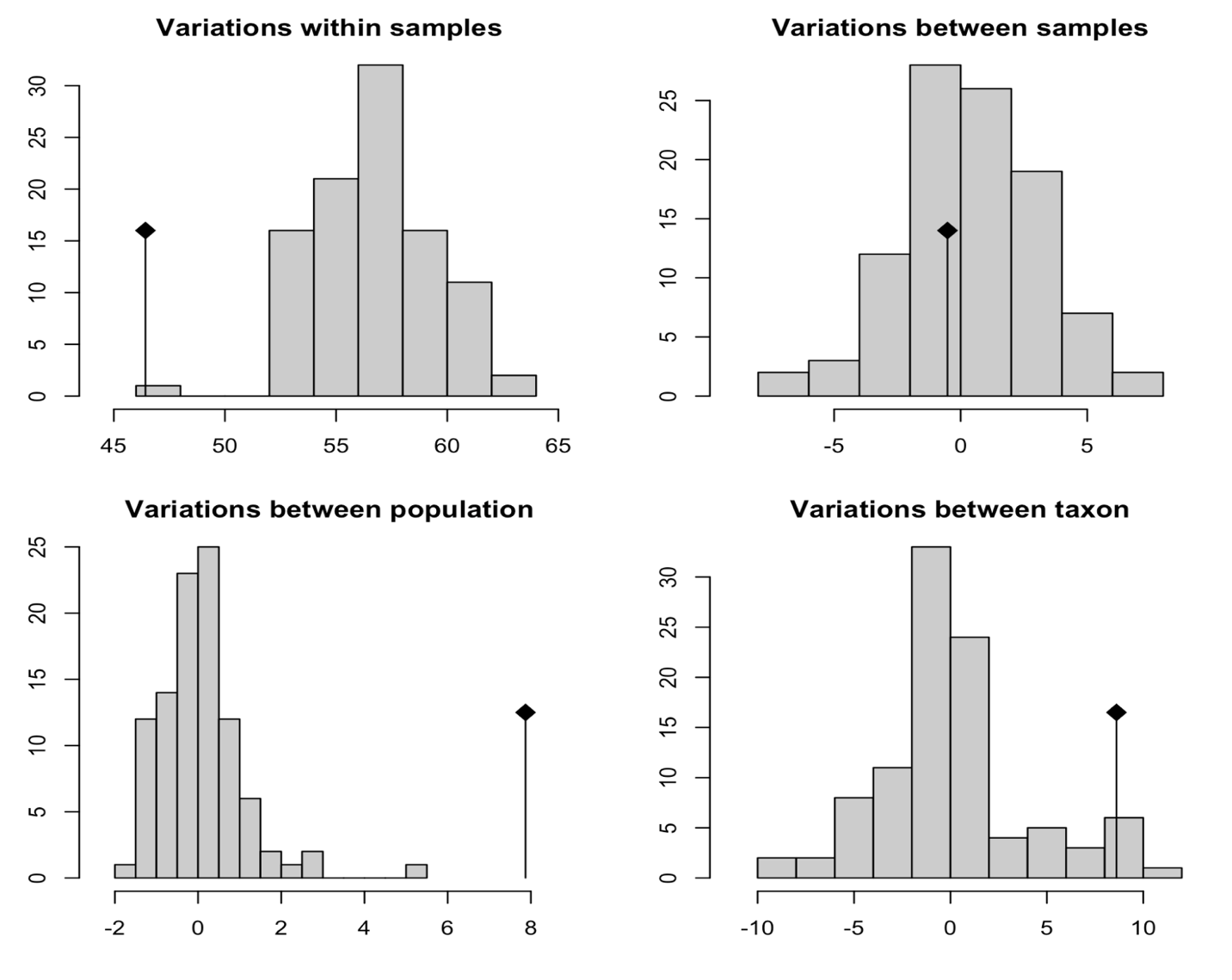

**Supplemental Table 3**: F_ST_ values for the *UCP1* gene region between savanna monkey populations, calculated using Nei’s (1987) distance.

|  | **East Africa** | **Zambia Kafue** | **Zambia-Chobe** | **FS North** | **FS South** | **KwaZulu-Natal** | **Eastern Cape** |
| --- | --- | --- | --- | --- | --- | --- | --- |
| **East Africa** | - | 0.2463 | 0.1008 | 0.1694 | 0.1828 | 0.2912 | 0.3805 |
| **Zambia Kafue** |  | - | 0.0989 | 0.2067 | 0.2972 | 0.3170 | 0.4479 |
| **Zambia-Chobe** |  |  | - | 0.0686 | 0.1141 | 0.2143 | 0.3315 |
| **FS North** |  |  |  | - | 0.0682 | 0.0643 | 0.1592 |
| **FS South** |  |  |  |  | - | 0.2379 | 0.3358 |
| **KwaZulu-Natal** |  |  |  |  |  | - | 0.0403 |
| **Eastern Cape** |  |  |  |  |  |  | - |

**Supplemental Figure 4**: Cross-entropy criterion values from *snmf* run in {LEA} for 1-20 potential clusters indicating 10 as the optimal number of ancestral populations representing the variation in the *UCP1* gene region in savanna monkeys.

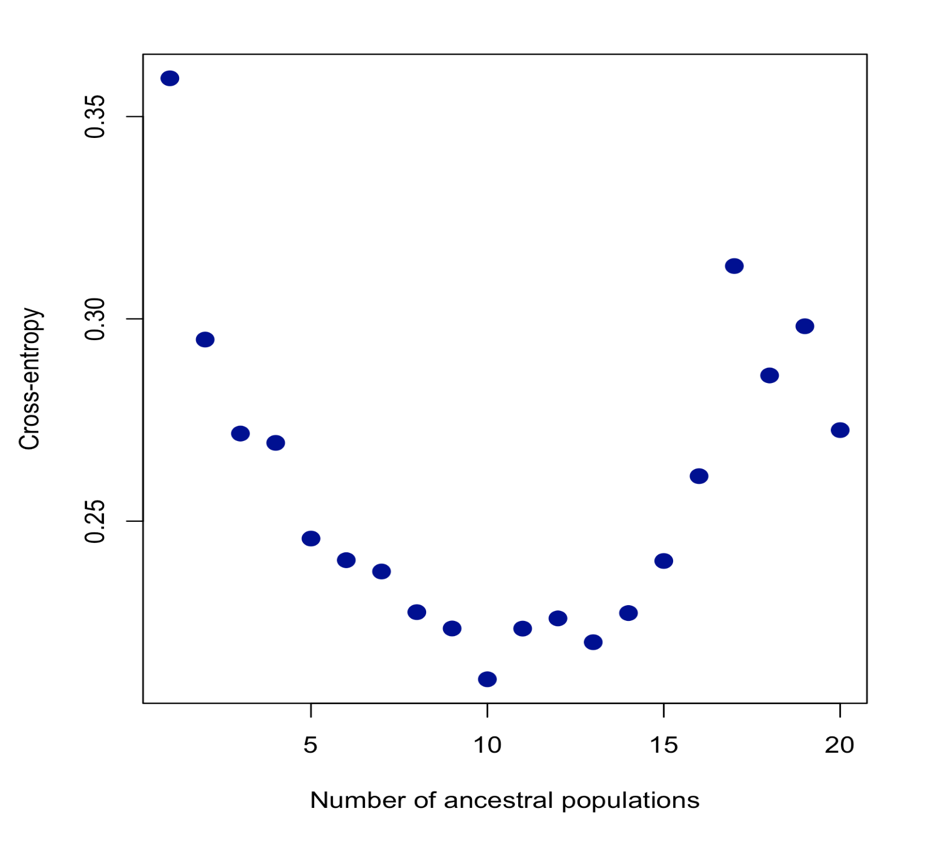

**Supplemental Figure 5**: Cluster assignments from {LEA} using k = 10 clusters to model the variation in the *UCP1* gene region in savanna monkeys. Clusters have been colored according to the taxon/population with the largest representation in the cluster, wherein shades of purple are most prominent in *C. p. hilgerti*, pink in *C. cynosuros*, and green in *C. p. pygerythrus*. Proportions of each per population can be seen in Figure 1A.

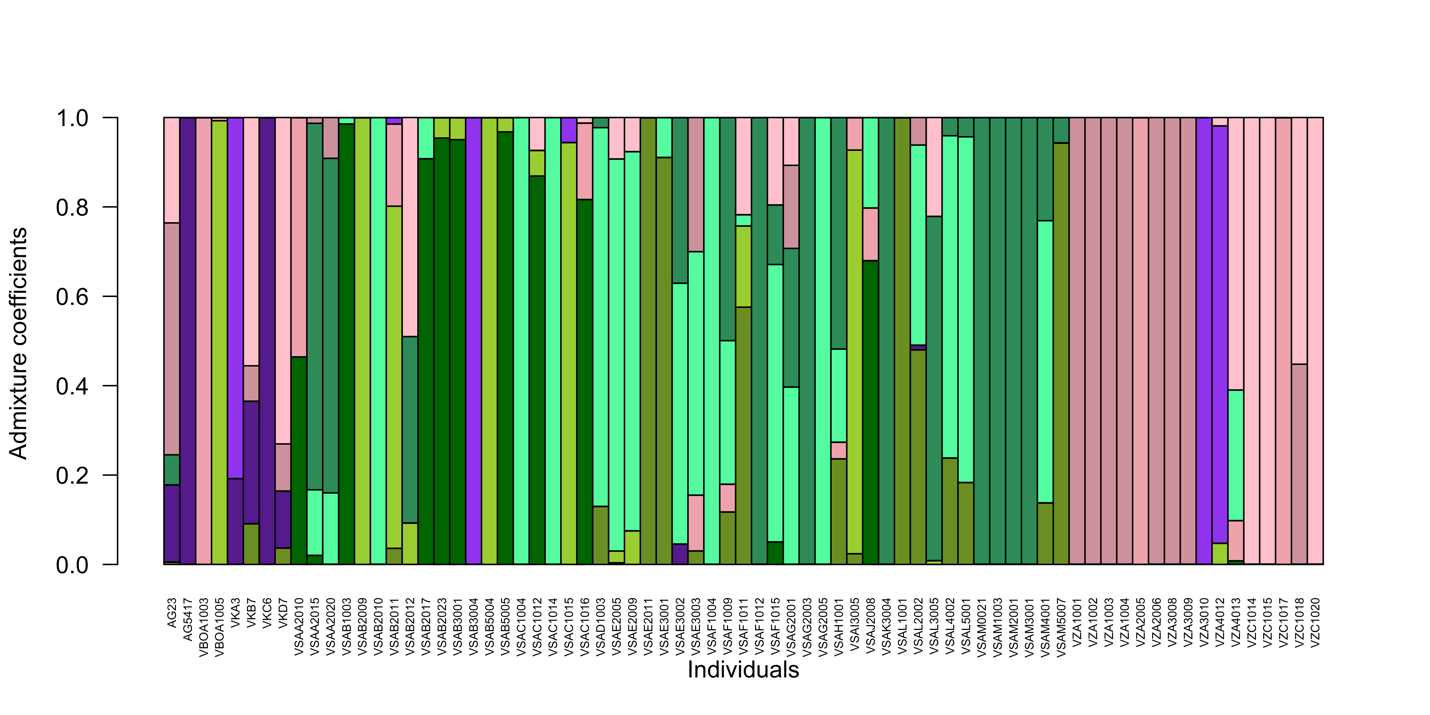

**Supplemental Table 4**: r^2^ LD block positions derived from the {gpart} package for the *UCP1* gene region in savanna monkeys.

| **LD Block** | **Chrom.** | **Start Pos.** | **End Pos.** | **Gene Region** |
| --- | --- | --- | --- | --- |
| B-01 | CAE7 | 87482499 | 87491914 | 3' Downstream |
| B-02 | CAE7 | 87492020 | 87493023 | 3' Downstream / 3’ UTR / Exon 6 / Intron 5-6 |
| B-03 | CAE7 | 87493125 | 87496052 | Intron 5-6 / Exon 5 / Intron 4-5 / Exon 4 / Intron 3-4 / Exon 3 |
| B-04 | CAE7 | 87497024 | 87497583 | Intron 2-3 |
| B-05 | CAE7 | 87497815 | 87498903 | Intron 2-3 |
| B-06 | CAE7 | 87499152 | 87500305 | Intron 2-3 |
| B-07 | CAE7 | 87500651 | 87501527 | Intron 2-3 / Exon 2 |
| B-08 | CAE7 | 87501699 | 87501756 | Intron 1-2 |
| B-09 | CAE7 | 87501834 | 87507475 | Intron 1-2 / Exon 1 / 5' UTR / 5' Upstream |
| B-10 | CAE7 | 87507524 | 87507706 | 5' Upstream |
| B-11 | CAE7 | 87508112 | 87508167 | 5' Upstream |
| B-12 | CAE7 | 87508195 | 87508268 | 5' Upstream |
| B-13 | CAE7 | 87508299 | 87508500 | 5' Upstream |
| B-14 | CAE7 | 87508541 | 87512621 | 5' Upstream |

**Supplemental Figure 6**: r^2^-based LD heatmap of the *UCP1* gene region of savanna monkeys using {gpart}. Positions for each block are illustrated but may also be found in Supplemental Table 4.

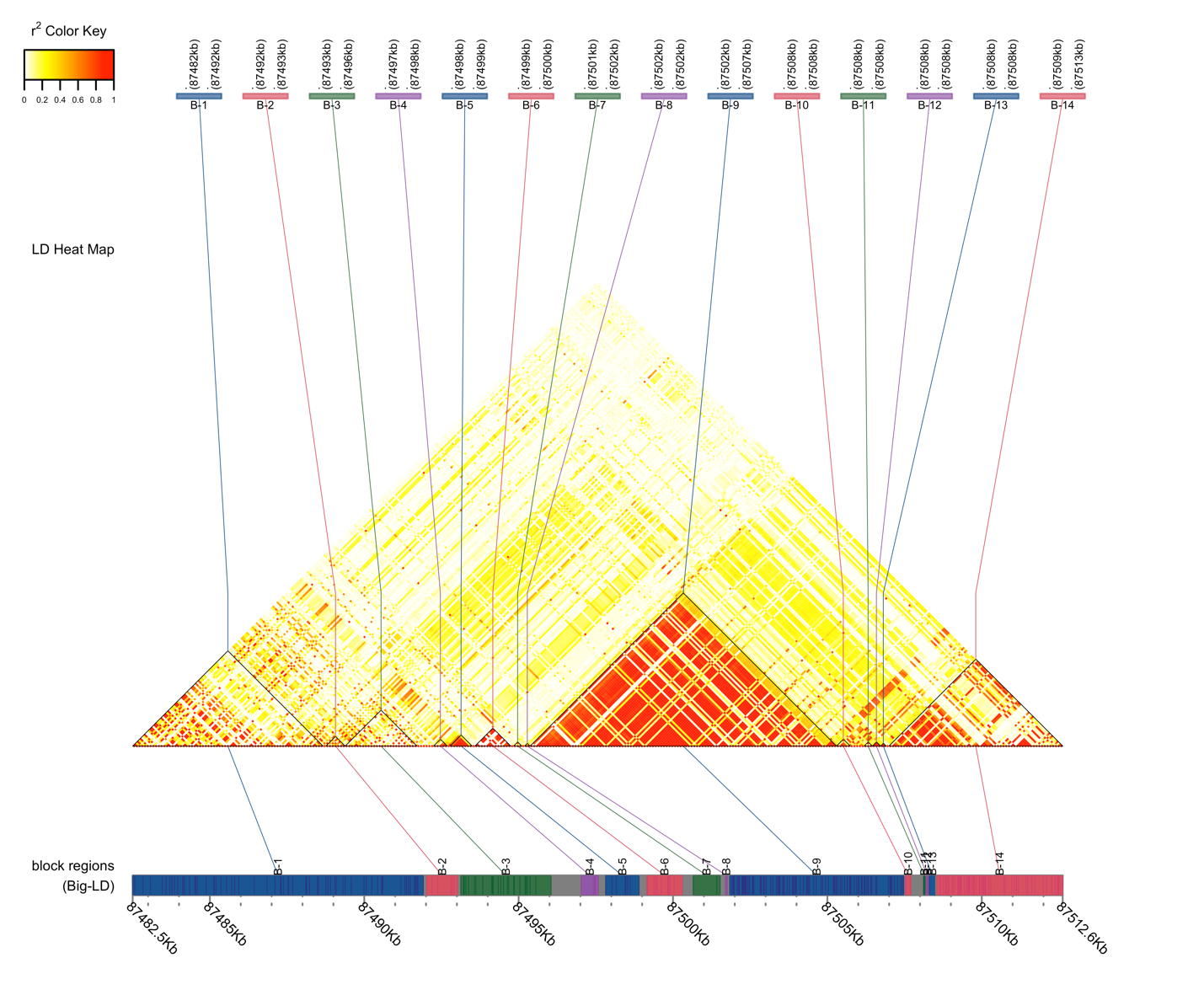

**Supplemental Figure 7**: LD (r^2^) of 10 loci of interest.

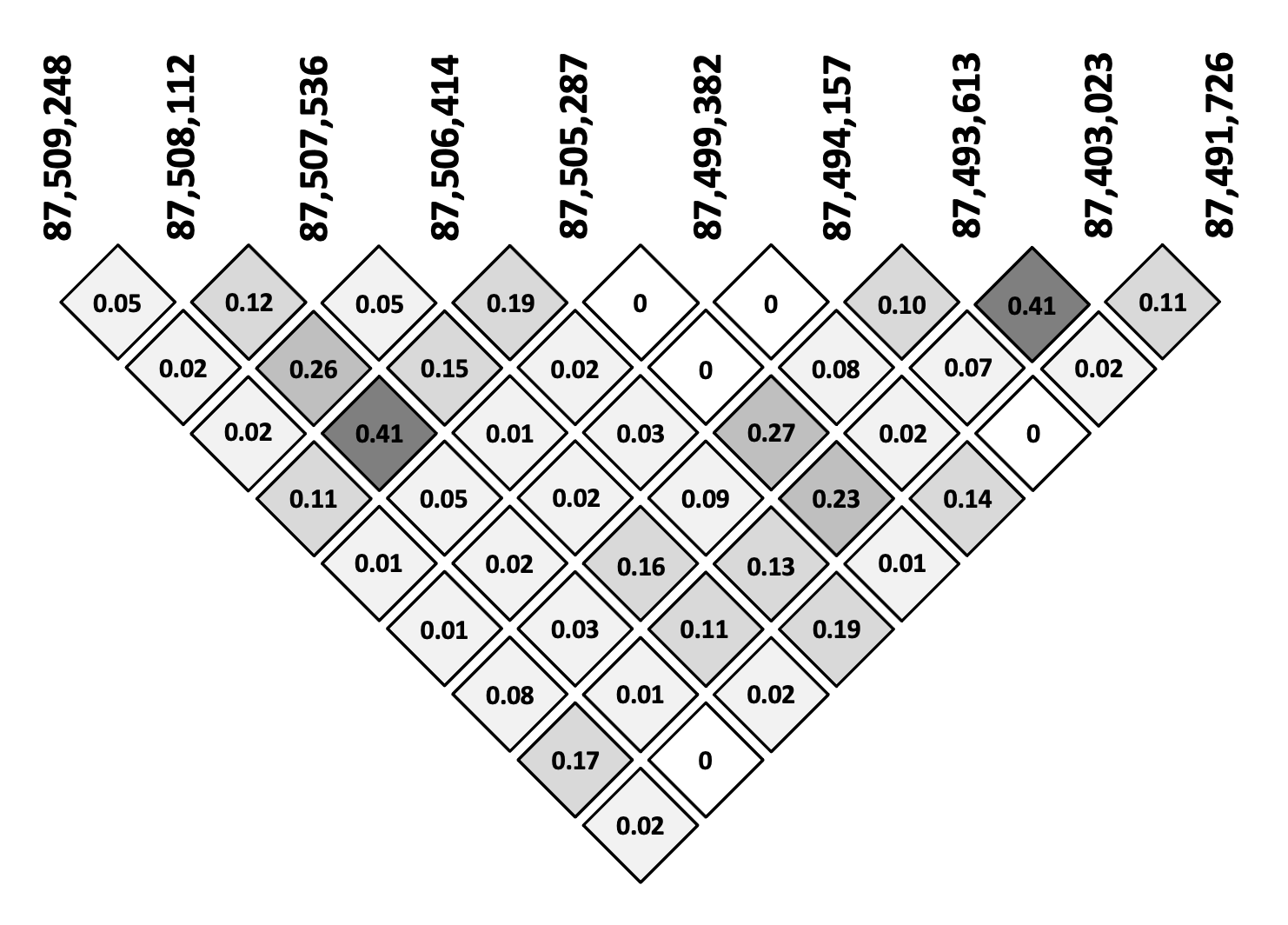

**Supplemental Table 6**: Sliding window Tajima’s D results in the *UCP1* region of savanna monkeys in a) the southern expansion, b) Free State North, c) Free State South, d) Eastern Cape, and e) KwaZulu-Natal using {vcftools} standard settings and a 500 bp window.

a) **Southern Expansion**

| **Chromosome** | **Start Position** | **N SNPs** | **Tajima's D** |
| --- | --- | --- | --- |
| CAE7 | 87482000 | 4 | -0.713316 |
| CAE7 | 87482500 | 5 | -0.631584 |
| CAE7 | 87483000 | 2 | 1.76628 |
| CAE7 | 87483500 | 7 | 0.625656 |
| CAE7 | 87484000 | 8 | -0.141022 |
| CAE7 | 87484500 | 7 | 0.194233 |
| CAE7 | 87485000 | 7 | 0.274367 |
| CAE7 | 87485500 | 6 | -0.0484465 |
| CAE7 | 87486000 | 7 | -0.885902 |
| CAE7 | 87486500 | 1 | 0.0338037 |
| CAE7 | 87487000 | 4 | 1.40705 |
| CAE7 | 87487500 | 5 | -0.604621 |
| CAE7 | 87488000 | 9 | -0.309112 |
| CAE7 | 87488500 | 6 | 0.371949 |
| CAE7 | 87489000 | 6 | 0.0155874 |
| CAE7 | 87489500 | 4 | -1.01056 |
| CAE7 | 87490000 | 7 | -0.969693 |
| CAE7 | 87490500 | 9 | -1.20137 |
| CAE7 | 87491000 | 6 | 0.465309 |
| CAE7 | 87491500 | 8 | 0.780017 |
| CAE7 | 87492000 | 6 | 0.609895 |
| CAE7 | 87492500 | 6 | 0.737963 |
| CAE7 | 87493000 | 8 | 1.00164 |
| CAE7 | 87493500 | 10 | -0.512656 |
| CAE7 | 87494000 | 10 | -0.243568 |
| CAE7 | 87494500 | 6 | 0.8703 |
| CAE7 | 87495000 | 7 | 0.765473 |
| CAE7 | 87495500 | 3 | -0.942327 |
| CAE7 | 87496000 | 4 | 1.04864 |
| CAE7 | 87496500 | 9 | -0.425907 |
| CAE7 | 87497000 | 9 | 0.303582 |
| **CAE7** | **87497500** | **4** | **2.04094** |
| **CAE7** | **87498000** | **5** | **2.32902** |
| CAE7 | 87498500 | 9 | -0.314772 |
| CAE7 | 87499000 | 17 | -1.37366 |
| CAE7 | 87499500 | 4 | -0.295701 |
| CAE7 | 87500000 | 10 | -1.63246 |
| CAE7 | 87500500 | 4 | -0.37668 |
| CAE7 | 87501000 | 3 | -0.597898 |
| CAE7 | 87501500 | 12 | 1.78182 |
| CAE7 | 87502000 | 21 | 1.3078 |
| CAE7 | 87502500 | 14 | 0.868846 |
| **CAE7** | **87503000** | **11** | **2.04155** |
| CAE7 | 87503500 | 13 | 1.38579 |
| CAE7 | 87504000 | 9 | 0.985579 |
| CAE7 | 87504500 | 9 | 0.0336818 |
| CAE7 | 87505000 | 13 | 0.950213 |
| CAE7 | 87505500 | 11 | 1.38946 |
| CAE7 | 87506000 | 18 | 0.912612 |
| CAE7 | 87506500 | 6 | 0.157761 |
| CAE7 | 87507000 | 14 | 1.5049 |
| CAE7 | 87507500 | 16 | 1.06294 |
| CAE7 | 87508000 | 11 | 1.56071 |
| **CAE7** | **87508500** | **11** | **2.84275** |
| CAE7 | 87509000 | 17 | 1.01075 |
| CAE7 | 87509500 | 18 | 1.86074 |
| CAE7 | 87510000 | 14 | 1.18892 |
| CAE7 | 87510500 | 6 | -0.882928 |
| CAE7 | 87511000 | 9 | -0.786648 |
| CAE7 | 87511500 | 5 | -0.123932 |
| CAE7 | 87512000 | 11 | 0.159842 |
| CAE7 | 87512500 | 4 | 0.86173 |

b) **Free State North**

| **Chromosome** | **Start Position** | **N SNPs** | **Tajima's D** |
| --- | --- | --- | --- |
| CAE7 | 87482000 | 2 | -0.527781 |
| CAE7 | 87482500 | 2 | -0.527781 |
| CAE7 | 87483000 | 2 | 0.895077 |
| CAE7 | 87483500 | 5 | 0.436778 |
| CAE7 | 87484000 | 4 | -0.0435264 |
| CAE7 | 87484500 | 4 | 0.321344 |
| CAE7 | 87485000 | 3 | 0.969734 |
| CAE7 | 87485500 | 3 | -0.227174 |
| CAE7 | 87486000 | 1 | 1.26176 |
| CAE7 | 87486500 | 1 | -1.16439 |
| CAE7 | 87487000 | 4 | 0.321344 |
| CAE7 | 87487500 | 3 | -0.227174 |
| CAE7 | 87488000 | 4 | 0.564591 |
| CAE7 | 87488500 | 4 | -0.0435264 |
| CAE7 | 87489000 | 4 | 0.280803 |
| CAE7 | 87489500 | 2 | -0.768572 |
| CAE7 | 87490000 | 4 | -0.16515 |
| CAE7 | 87490500 | 5 | -1.003 |
| CAE7 | 87491000 | 5 | -0.0202956 |
| CAE7 | 87491500 | 7 | -0.770801 |
| CAE7 | 87492000 | 4 | 0.388912 |
| CAE7 | 87492500 | 3 | 1.50169 |
| CAE7 | 87493000 | 6 | 0.403337 |
| CAE7 | 87493500 | 3 | 0.37128 |
| CAE7 | 87494000 | 5 | 0.311083 |
| CAE7 | 87494500 | 4 | 0.145666 |
| CAE7 | 87495000 | 3 | 1.0861 |
| CAE7 | 87495500 | 1 | -1.16439 |
| CAE7 | 87496000 | 2 | 0.0632525 |
| CAE7 | 87496500 | 3 | 0.304785 |
| CAE7 | 87497000 | 6 | 0.60169 |
| CAE7 | 87497500 | 4 | 0.618646 |
| CAE7 | 87498000 | 4 | 1.64569 |
| CAE7 | 87498500 | 4 | 0.348371 |
| CAE7 | 87499000 | 8 | -1.23852 |
| CAE7 | 87499500 | 3 | -0.626144 |
| CAE7 | 87500000 | 3 | -1.1581 |
| CAE7 | 87500500 | 2 | 0.172703 |
| CAE7 | 87501000 | 2 | -0.286989 |
| CAE7 | 87501500 | 10 | 1.11295 |
| CAE7 | 87502000 | 19 | 0.782163 |
| CAE7 | 87502500 | 10 | 0.773413 |
| CAE7 | 87503000 | 10 | 1.17172 |
| CAE7 | 87503500 | 12 | 1.62173 |
| CAE7 | 87504000 | 7 | 1.0359 |
| CAE7 | 87504500 | 5 | 1.22523 |
| CAE7 | 87505000 | 9 | 1.29855 |
| CAE7 | 87505500 | 9 | 1.54827 |
| CAE7 | 87506000 | 13 | 1.22325 |
| CAE7 | 87506500 | 3 | 1.46845 |
| **CAE7** | **87507000** | **10** | **2.1838** |
| **CAE7** | **87507500** | **4** | **2.41597** |
| **CAE7** | **87508000** | **5** | **2.01368** |
| **CAE7** | **87508500** | **9** | **2.64702** |
| **CAE7** | **87509000** | **9** | **2.07624** |
| **CAE7** | **87509500** | **10** | **2.7127** |
| CAE7 | 87510000 | 8 | 0.342652 |
| **CAE7** | **87510500** | **2** | **-1.51284** |
| CAE7 | 87511000 | 1 | -1.16439 |
| CAE7 | 87511500 | 2 | 0.938857 |
| CAE7 | 87512000 | 4 | 1.56461 |
| **CAE7** | **87512500** | **2** | **1.98958** |

c) **Free State South**

| **Chromosome** | **Start Position** | **N SNPs** | **Tajima's D** |
| --- | --- | --- | --- |
| CAE7 | 87482000 | 1 | 0.895275 |
| CAE7 | 87482500 | 2 | 1.1667 |
| CAE7 | 87483000 | 2 | 0.469505 |
| CAE7 | 87483500 | 5 | 0.686291 |
| CAE7 | 87484000 | 4 | 0.825702 |
| CAE7 | 87484500 | 4 | 1.01405 |
| CAE7 | 87485000 | 3 | 0.801325 |
| CAE7 | 87485500 | 3 | 1.00541 |
| CAE7 | 87486000 | 1 | -0.174717 |
| CAE7 | 87486500 | 1 | 0.895275 |
| CAE7 | 87487000 | 4 | 0.770306 |
| CAE7 | 87487500 | 3 | 0.896566 |
| CAE7 | 87488000 | 5 | 0.301676 |
| CAE7 | 87488500 | 4 | 1.71204 |
| CAE7 | 87489000 | 2 | 1.54211 |
| CAE7 | 87489500 | 0 | nan |
| CAE7 | 87490000 | 3 | -1.21235 |
| CAE7 | 87490500 | 2 | -1.51481 |
| CAE7 | 87491000 | 4 | 1.32427 |
| **CAE7** | **87491500** | **6** | **2.74507** |
| CAE7 | 87492000 | 4 | 1.11376 |
| CAE7 | 87492500 | 3 | 1.08705 |
| CAE7 | 87493000 | 6 | 0.837813 |
| **CAE7** | **87493500** | **3** | **1.94422** |
| CAE7 | 87494000 | 5 | 0.0952977 |
| CAE7 | 87494500 | 4 | 0.659513 |
| CAE7 | 87495000 | 3 | 0.760507 |
| CAE7 | 87495500 | 1 | 0.895275 |
| CAE7 | 87496000 | 2 | 1.54211 |
| CAE7 | 87496500 | 2 | -1.17515 |
| CAE7 | 87497000 | 7 | -0.263883 |
| CAE7 | 87497500 | 4 | -0.681074 |
| CAE7 | 87498000 | 4 | -0.282221 |
| CAE7 | 87498500 | 5 | -1.13359 |
| CAE7 | 87499000 | 7 | -0.884312 |
| CAE7 | 87499500 | 3 | -1.47087 |
| CAE7 | 87500000 | 3 | -0.382391 |
| CAE7 | 87500500 | 2 | -1.17515 |
| CAE7 | 87501000 | 1 | -1.1624 |
| **CAE7** | **87501500** | **9** | **2.34309** |
| **CAE7** | **87502000** | **18** | **2.91988** |
| **CAE7** | **87502500** | **10** | **2.70861** |
| **CAE7** | **87503000** | **10** | **2.27808** |
| **CAE7** | **87503500** | **11** | **3.10733** |
| **CAE7** | **87504000** | **8** | **2.24093** |
| **CAE7** | **87504500** | **5** | **2.65627** |
| **CAE7** | **87505000** | **8** | **2.57118** |
| **CAE7** | **87505500** | **9** | **2.94843** |
| **CAE7** | **87506000** | **12** | **2.39408** |
| **CAE7** | **87506500** | **3** | **2.09389** |
| **CAE7** | **87507000** | **10** | **2.61712** |
| CAE7 | 87507500 | 5 | 1.60561 |
| **CAE7** | **87508000** | **4** | **2.15521** |
| **CAE7** | **87508500** | **9** | **2.24906** |
| **CAE7** | **87509000** | **8** | **2.58414** |
| **CAE7** | **87509500** | **10** | **2.61712** |
| CAE7 | 87510000 | 10 | -0.0144359 |
| CAE7 | 87510500 | 0 | nan |
| CAE7 | 87511000 | 0 | nan |
| CAE7 | 87511500 | 1 | 1.33425 |
| CAE7 | 87512000 | 4 | 1.35751 |
| CAE7 | 87512500 | 2 | 1.73875 |

d) **Eastern Cape**

| **Chromosome** | **Start Position** | **N SNPs** | **Tajima's D** |
| --- | --- | --- | --- |
| CAE7 | 87482000 | 1 | -1.15933 |
| CAE7 | 87482500 | 2 | -1.51469 |
| CAE7 | 87483000 | 2 | -1.20229 |
| CAE7 | 87483500 | 3 | -1.49431 |
| CAE7 | 87484000 | 4 | -1.88381 |
| CAE7 | 87484500 | 4 | -1.68955 |
| CAE7 | 87485000 | 3 | -1.49431 |
| CAE7 | 87485500 | 3 | -1.73253 |
| CAE7 | 87486000 | 2 | -1.20229 |
| CAE7 | 87486500 | 1 | -1.15933 |
| CAE7 | 87487000 | 4 | -1.68955 |
| CAE7 | 87487500 | 2 | -1.51469 |
| CAE7 | 87488000 | 5 | -0.294444 |
| CAE7 | 87488500 | 3 | -1.73253 |
| CAE7 | 87489000 | 2 | -1.51469 |
| CAE7 | 87489500 | 0 | nan |
| CAE7 | 87490000 | 3 | -0.0876744 |
| CAE7 | 87490500 | 0 | nan |
| CAE7 | 87491000 | 3 | -0.643523 |
| CAE7 | 87491500 | 1 | -1.15933 |
| CAE7 | 87492000 | 2 | -1.51469 |
| CAE7 | 87492500 | 3 | -1.49431 |
| CAE7 | 87493000 | 3 | -1.01787 |
| CAE7 | 87493500 | 3 | -1.49431 |
| CAE7 | 87494000 | 4 | -1.49528 |
| CAE7 | 87494500 | 3 | -1.49431 |
| CAE7 | 87495000 | 1 | -0.681114 |
| CAE7 | 87495500 | 2 | -1.51469 |
| CAE7 | 87496000 | 2 | -1.51469 |
| CAE7 | 87496500 | 1 | -1.15933 |
| CAE7 | 87497000 | 3 | -0.586804 |
| CAE7 | 87497500 | 3 | -0.371271 |
| CAE7 | 87498000 | 4 | -0.403689 |
| CAE7 | 87498500 | 3 | 0.36608 |
| CAE7 | 87499000 | 2 | -1.51469 |
| CAE7 | 87499500 | 1 | -1.15933 |
| CAE7 | 87500000 | 0 | nan |
| CAE7 | 87500500 | 1 | -0.248438 |
| CAE7 | 87501000 | 0 | nan |
| CAE7 | 87501500 | 9 | -0.488731 |
| CAE7 | 87502000 | 1 | 1.39118 |
| CAE7 | 87502500 | 1 | 1.39118 |
| **CAE7** | **87503000** | **3** | **2.079** |
| CAE7 | 87503500 | 0 | nan |
| CAE7 | 87504000 | 1 | 1.39118 |
| CAE7 | 87504500 | 0 | nan |
| CAE7 | 87505000 | 3 | 1.8975 |
| CAE7 | 87505500 | 3 | -0.462022 |
| **CAE7** | **87506000** | **5** | **2.28551** |
| CAE7 | 87506500 | 1 | 1.39118 |
| **CAE7** | **87507000** | **5** | **2.3404** |
| CAE7 | 87507500 | 7 | 0.468991 |
| **CAE7** | **87508000** | **4** | **2.19578** |
| **CAE7** | **87508500** | **9** | **2.60567** |
| CAE7 | 87509000 | 10 | 1.69688 |
| CAE7 | 87509500 | 13 | 1.73736 |
| **CAE7** | **87510000** | **10** | **2.25212** |
| CAE7 | 87510500 | 1 | -1.15933 |
| **CAE7** | **87511000** | **5** | **-1.99611** |
| CAE7 | 87511500 | 3 | 0.502206 |
| CAE7 | 87512000 | 8 | 0.0364814 |
| CAE7 | 87512500 | 3 | 0.967304 |

e) **KwaZulu-Natal**

| **Chromosome** | **Start Position** | **N SNPs** | **Tajima's D** |
| --- | --- | --- | --- |
| CAE7 | 87482000 | 1 | -0.681114 |
| CAE7 | 87482500 | 3 | -1.25609 |
| CAE7 | 87483000 | 2 | 0.35972 |
| CAE7 | 87483500 | 4 | -0.329683 |
| CAE7 | 87484000 | 4 | -0.77372 |
| CAE7 | 87484500 | 4 | -0.329683 |
| CAE7 | 87485000 | 4 | -0.329683 |
| CAE7 | 87485500 | 2 | -0.889887 |
| CAE7 | 87486000 | 1 | 1.23177 |
| CAE7 | 87486500 | 1 | -0.681114 |
| CAE7 | 87487000 | 4 | -0.329683 |
| CAE7 | 87487500 | 2 | -0.354341 |
| CAE7 | 87488000 | 5 | -0.906104 |
| CAE7 | 87488500 | 4 | -1.51378 |
| CAE7 | 87489000 | 3 | -0.643523 |
| CAE7 | 87489500 | 0 | nan |
| CAE7 | 87490000 | 3 | -0.269176 |
| CAE7 | 87490500 | 0 | nan |
| CAE7 | 87491000 | 2 | 0.746504 |
| CAE7 | 87491500 | 1 | 1.23177 |
| CAE7 | 87492000 | 2 | 0.463854 |
| CAE7 | 87492500 | 3 | 0.80849 |
| **CAE7** | **87493000** | **3** | **2.19244** |
| CAE7 | 87493500 | 1 | 1.50504 |
| CAE7 | 87494000 | 3 | 1.10343 |
| CAE7 | 87494500 | 2 | 1.49032 |
| CAE7 | 87495000 | 2 | 0.0621947 |
| CAE7 | 87495500 | 0 | nan |
| CAE7 | 87496000 | 3 | -0.371271 |
| CAE7 | 87496500 | 3 | 1.17149 |
| CAE7 | 87497000 | 4 | 1.15044 |
| CAE7 | 87497500 | 3 | 1.63659 |
| **CAE7** | **87498000** | **4** | **2.26053** |
| CAE7 | 87498500 | 3 | 1.77272 |
| CAE7 | 87499000 | 1 | -0.248438 |
| CAE7 | 87499500 | 1 | 0.480279 |
| CAE7 | 87500000 | 1 | -0.681114 |
| CAE7 | 87500500 | 1 | 1.39118 |
| CAE7 | 87501000 | 1 | 0.480279 |
| CAE7 | 87501500 | 9 | -0.321199 |
| CAE7 | 87502000 | 7 | -1.46418 |
| CAE7 | 87502500 | 2 | 0.865514 |
| CAE7 | 87503000 | 3 | 1.84078 |
| CAE7 | 87503500 | 0 | nan |
| CAE7 | 87504000 | 1 | 1.23177 |
| CAE7 | 87504500 | 0 | nan |
| CAE7 | 87505000 | 3 | 1.23956 |
| CAE7 | 87505500 | 3 | 0.150546 |
| CAE7 | 87506000 | 6 | 1.36768 |
| CAE7 | 87506500 | 1 | 1.50504 |
| CAE7 | 87507000 | 5 | 1.39154 |
| CAE7 | 87507500 | 13 | 0.217272 |
| **CAE7** | **87508000** | **7** | **1.94303** |
| **CAE7** | **87508500** | **10** | **2.29275** |
| CAE7 | 87509000 | 11 | 1.81212 |
| CAE7 | 87509500 | 14 | 1.6057 |
| CAE7 | 87510000 | 12 | 0.982841 |
| CAE7 | 87510500 | 4 | -0.551701 |
| CAE7 | 87511000 | 6 | -1.59216 |
| CAE7 | 87511500 | 3 | 0.445487 |
| CAE7 | 87512000 | 7 | 0.0581931 |
| **CAE7** | **87512500** | **2** | **2.01099** |

**Supplemental Table 7**: Significant p_iHS_ values for the *UCP1* region in savanna monkeys. Although the typical convention is to use a significance threshold of p_iHS_ = 2 (corresponding roughly to p = 0.01), we chose to investigate SNPs with p_iHS_ around 1.3 and above (corresponding to a p = 0.05).

| **Position** | **piHS** |
| --- | --- |
| 87491726 | 1.280563 |
| 87493023 | 1.287135 |
| 87493613 | 1.341058 |
| 87494157 | 1.850070 |
| 87496971 | 1.701703 |
| 87499382 | 1.566911 |
| 87500036 | 1.278484 |
| 87503019 | 1.474165 |
| 87505287 | 1.721821 |
| **87506414** | **2.837496** |
| 87506571 | 1.354139 |
| 87507536 | 1.838767 |
| **87508112** | **2.006234** |
| **87508167** | **2.290936** |
| **87509248** | **2.714808** |
| **87509754** | **2.262098** |
| 87510003 | 1.282514 |

**Supplemental Figure 8**: Integrated haplotype score (iHS) values across the *UCP1* gene region in savanna monkeys. The shaded region indicates the most likely region to be undergoing a positive selective sweep. This region corresponds to the exon 1 / 5’UTR / 5’ upstream region.

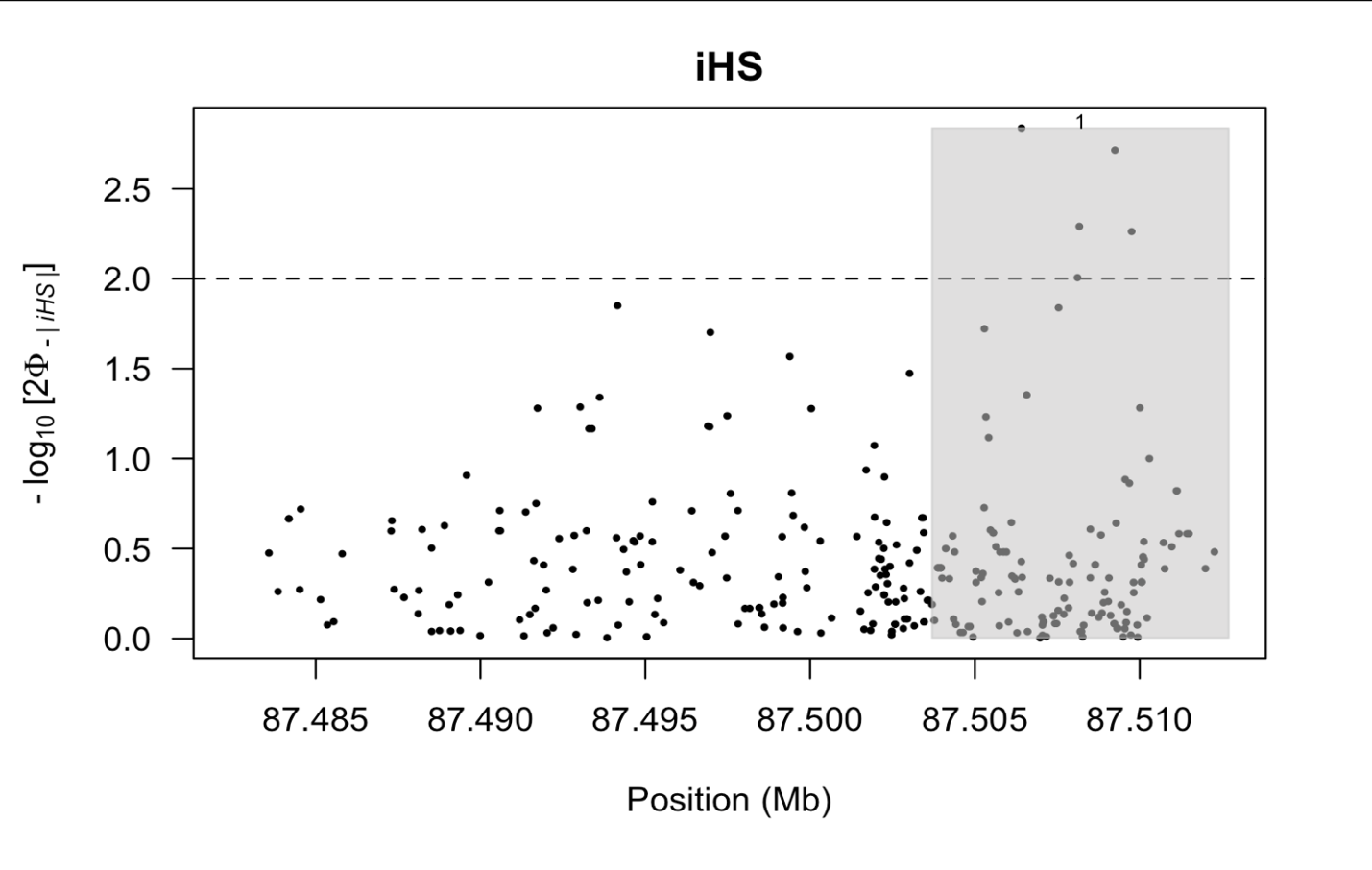

**Supplemental Figure 9**: Extended haplotype homozygosity (EHH) in 10 SNPs of interest (a-j) potentially having undergone recent positive selective sweeps in the *UCP1* gene region of savanna monkeys using i) diagrams of EHH, and ii) bifurcation diagrams illustrating sequence divergence (or breaking) of haplotypes branching from the derived (in yellow) and blue (ancestral) alleles at the SNP of interest.

a) **87491726**

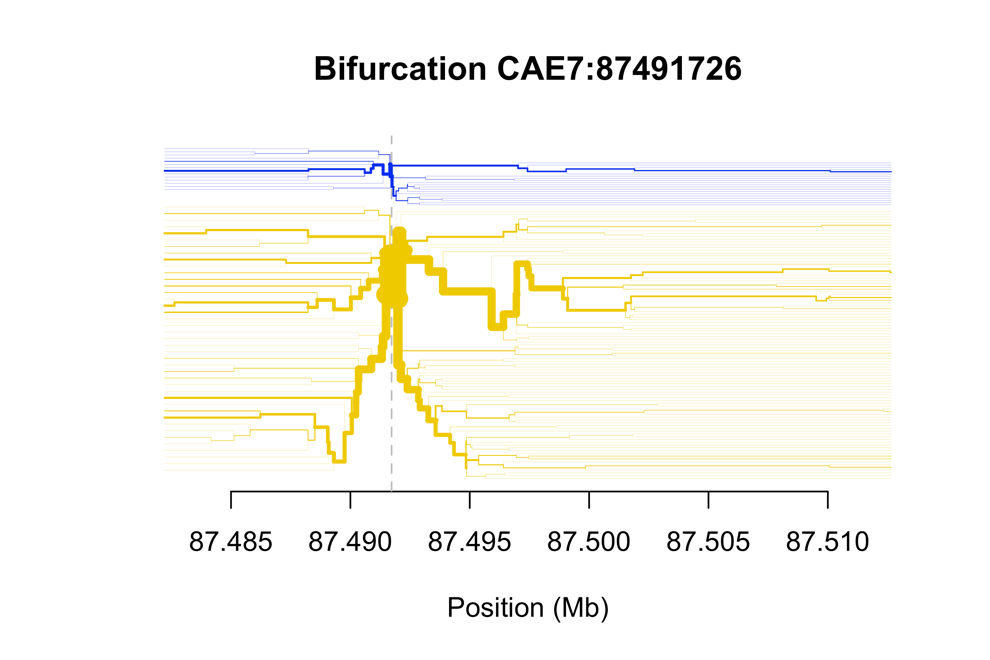

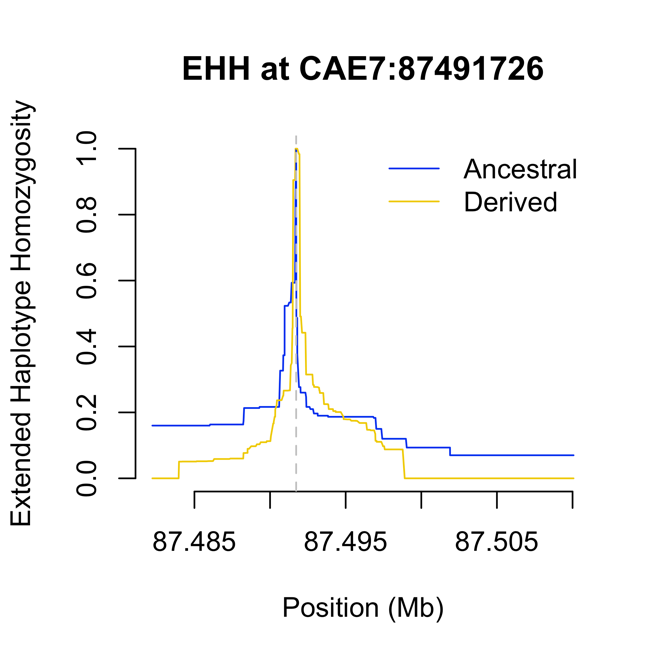

b) **87493023**

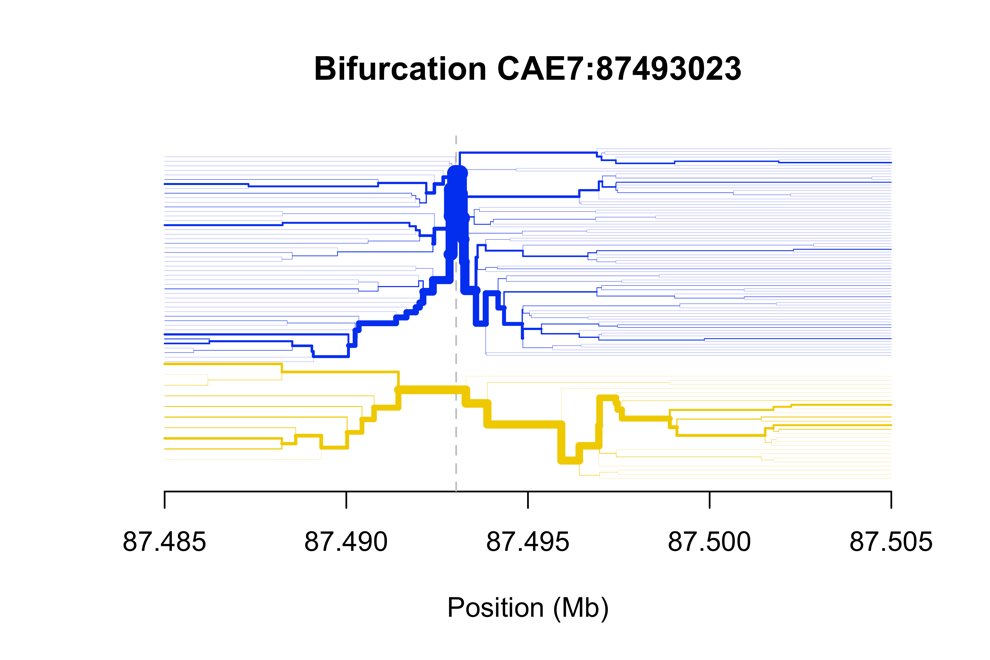

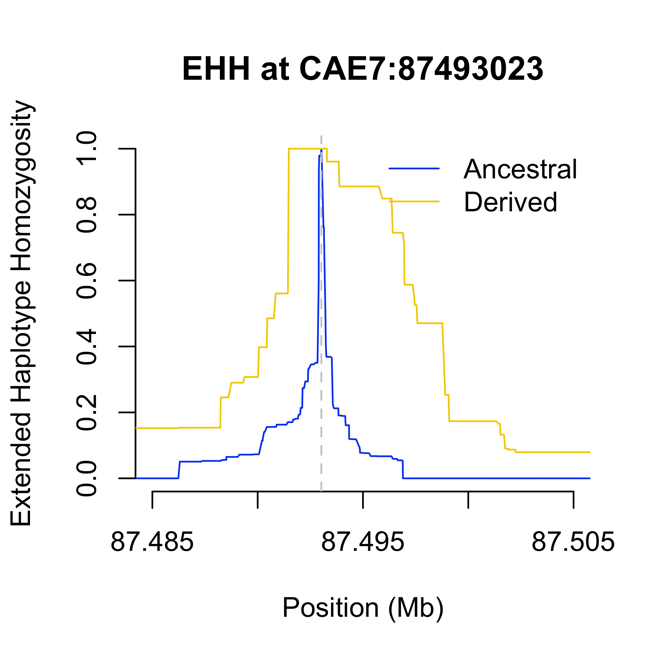

c) **87493613**

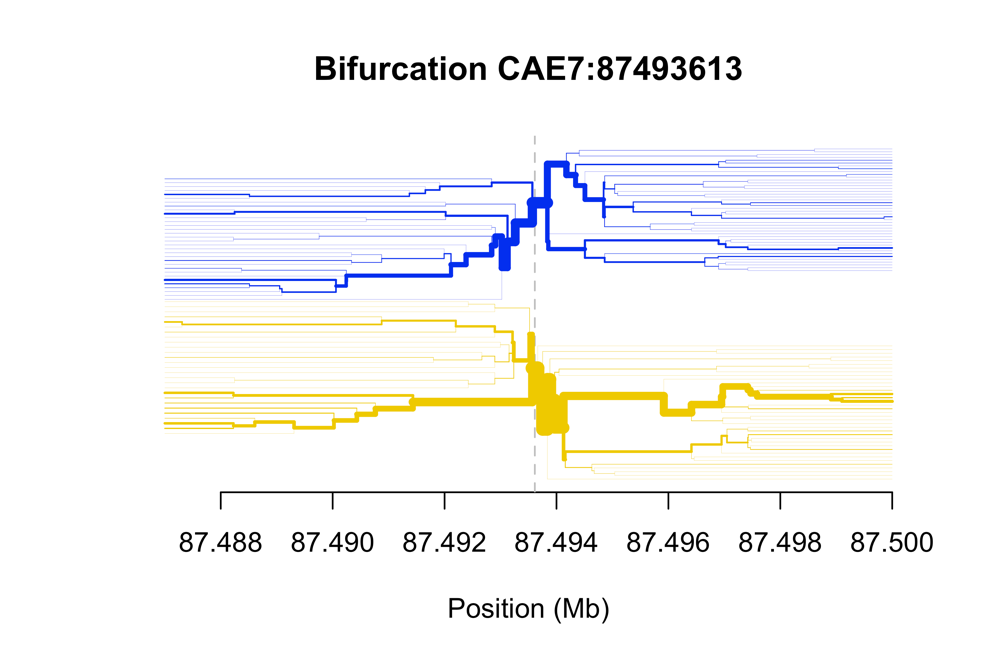

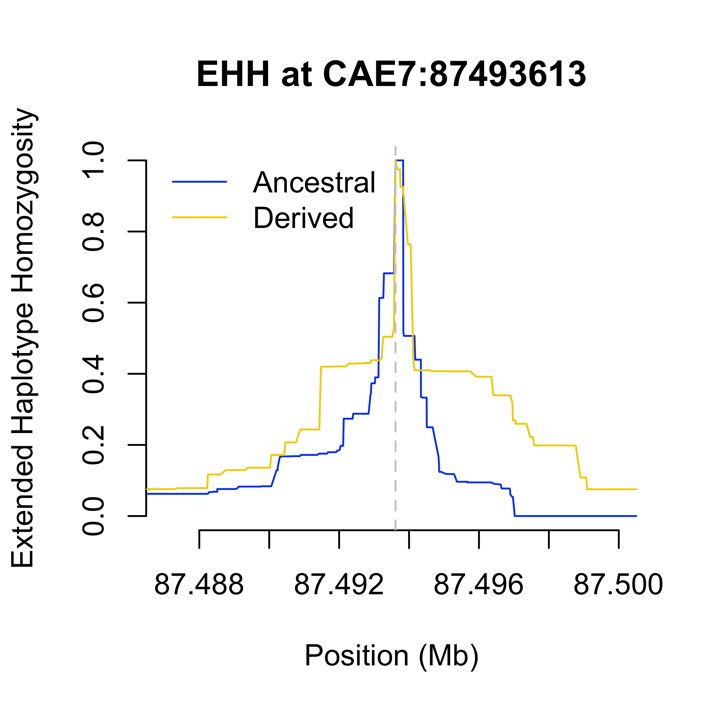

d) **87494157**

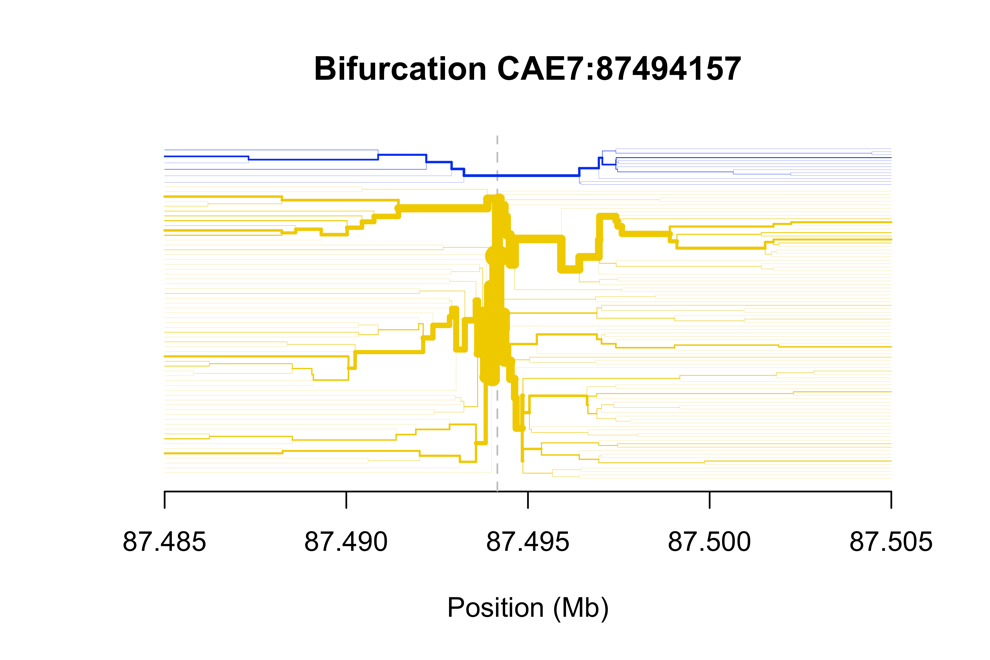

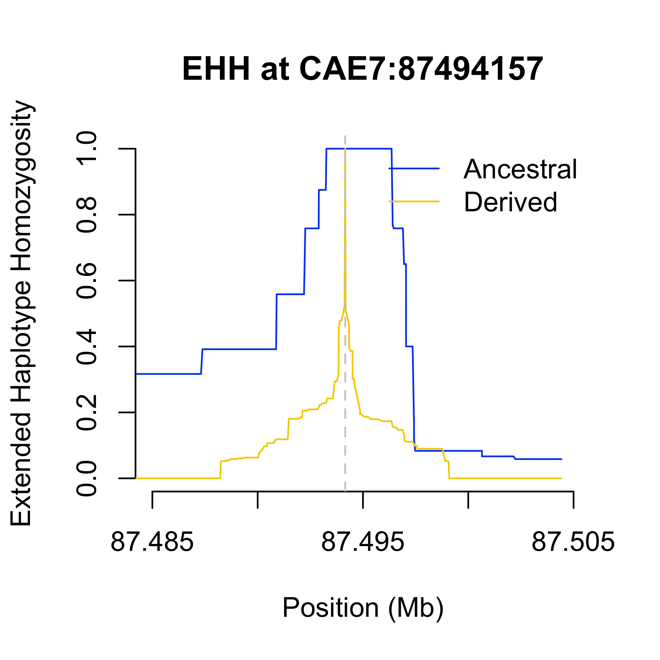

e) **87499382**

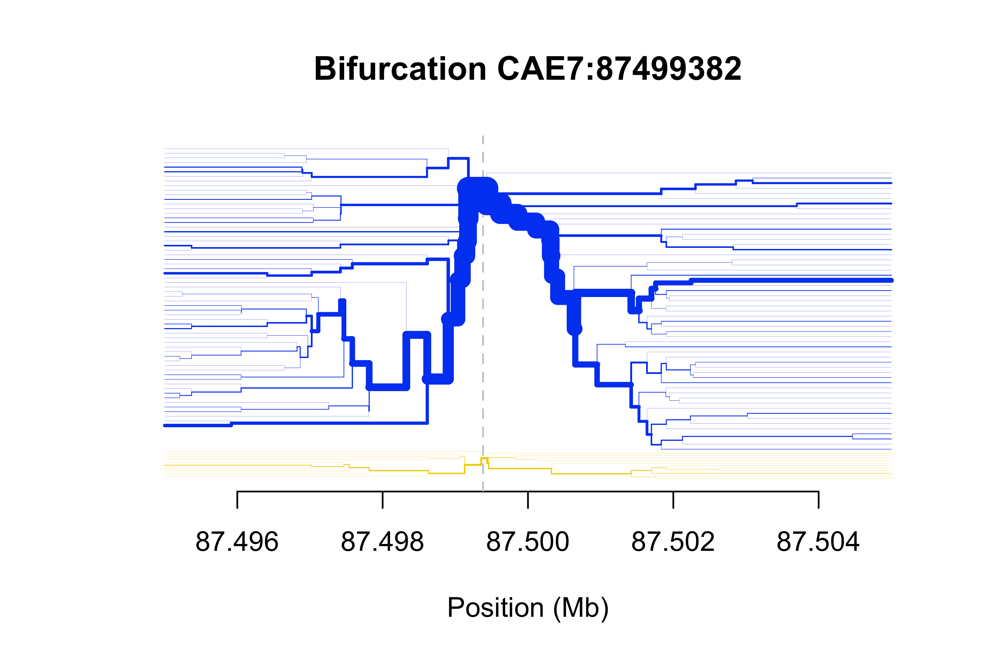

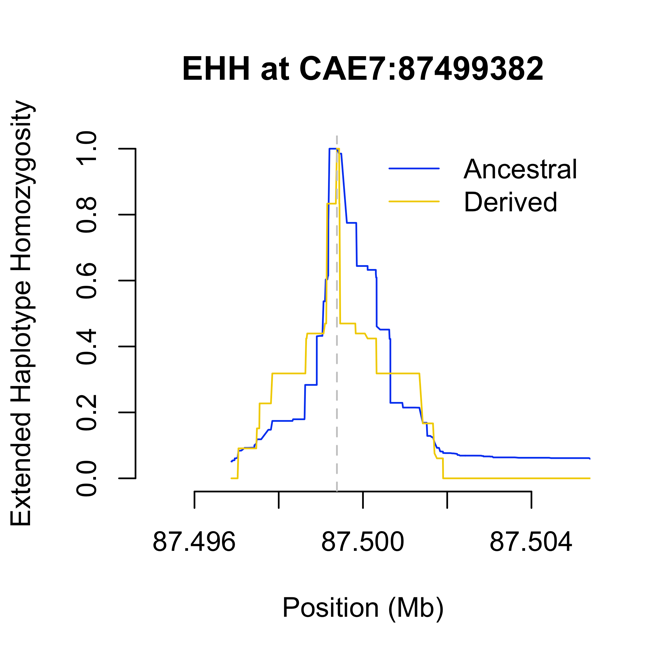

f) **87505287**

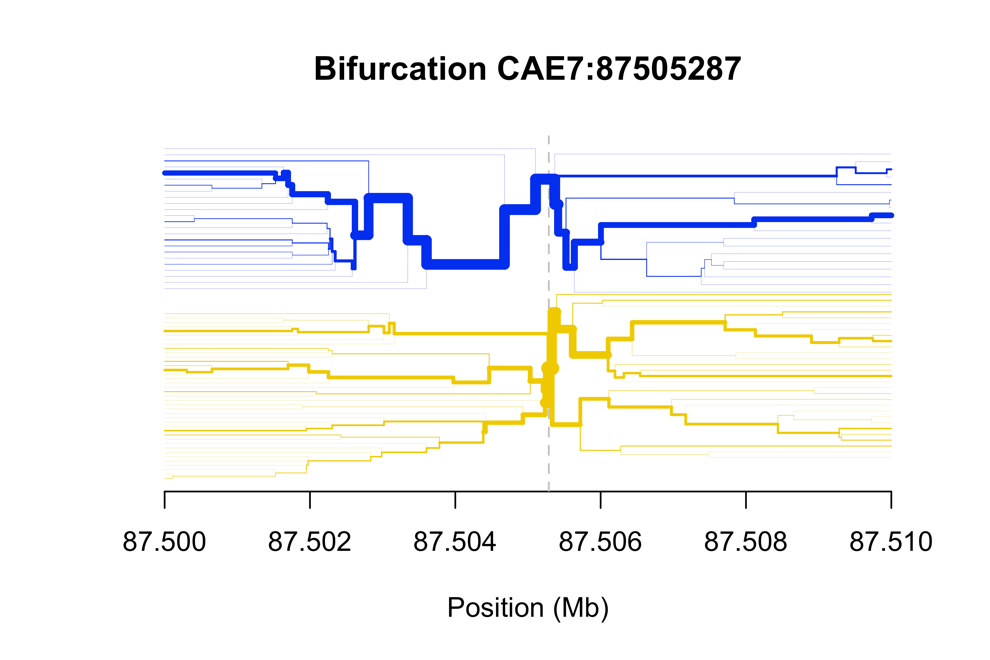

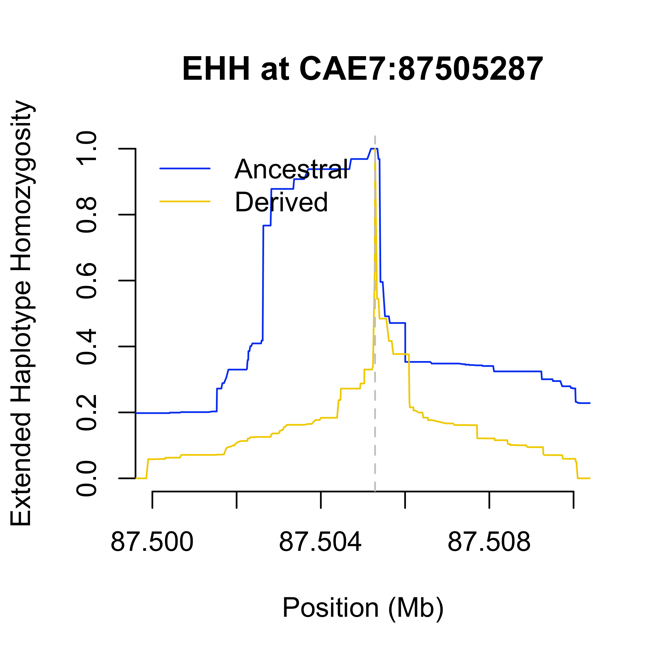

g) **87506414**

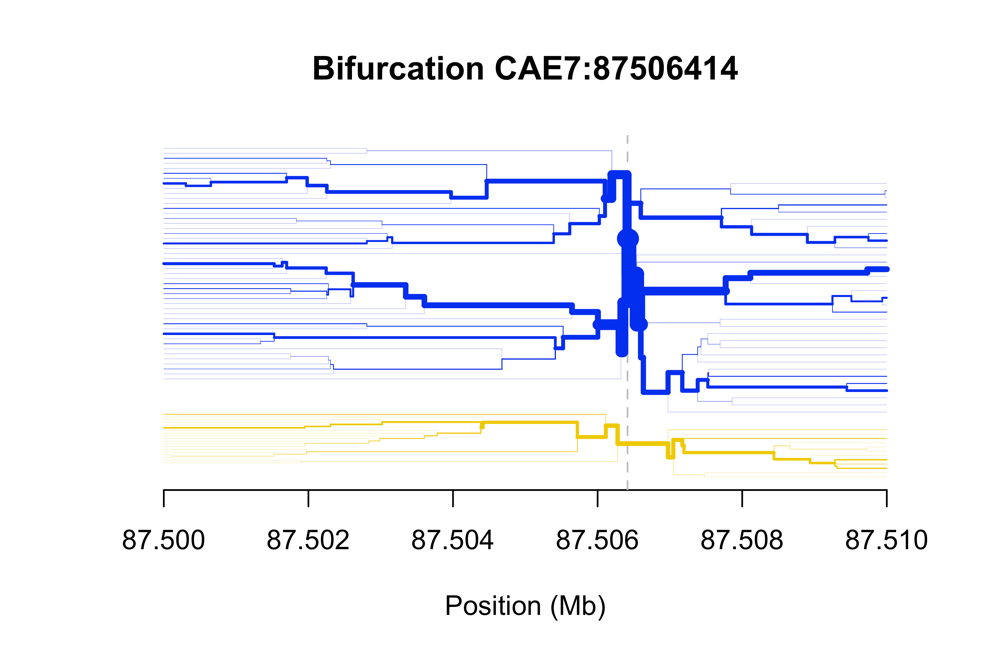

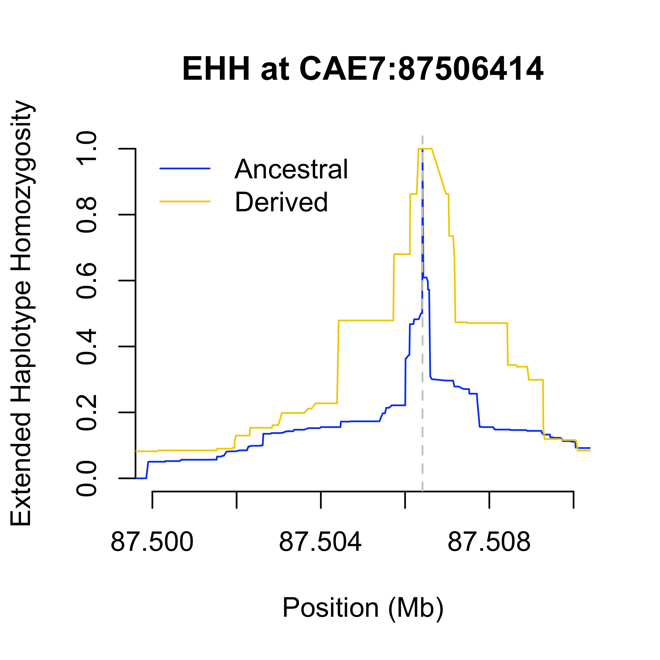

h) **87507536**

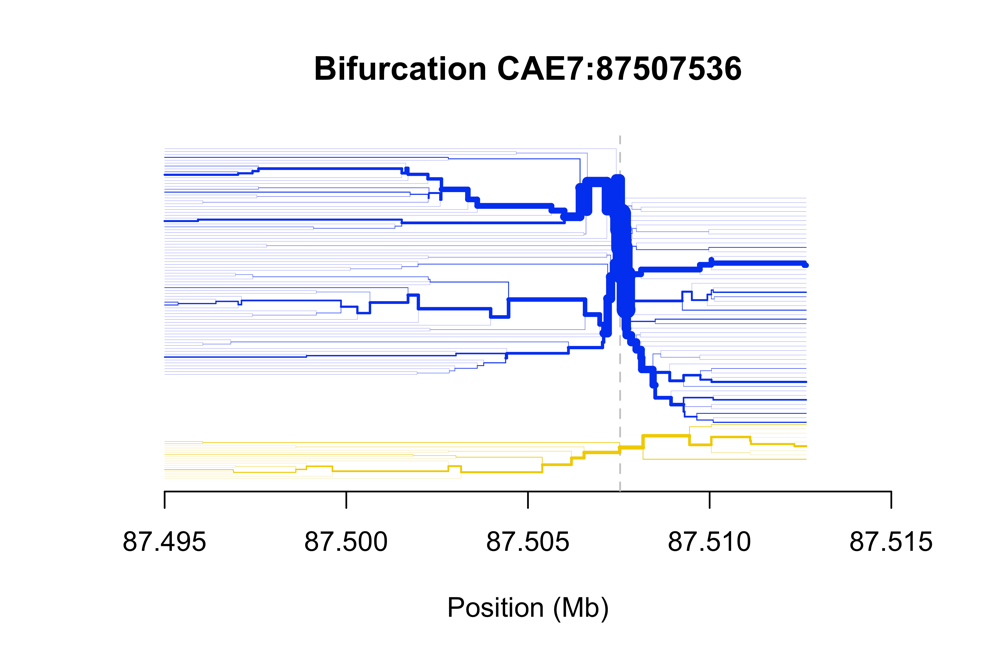

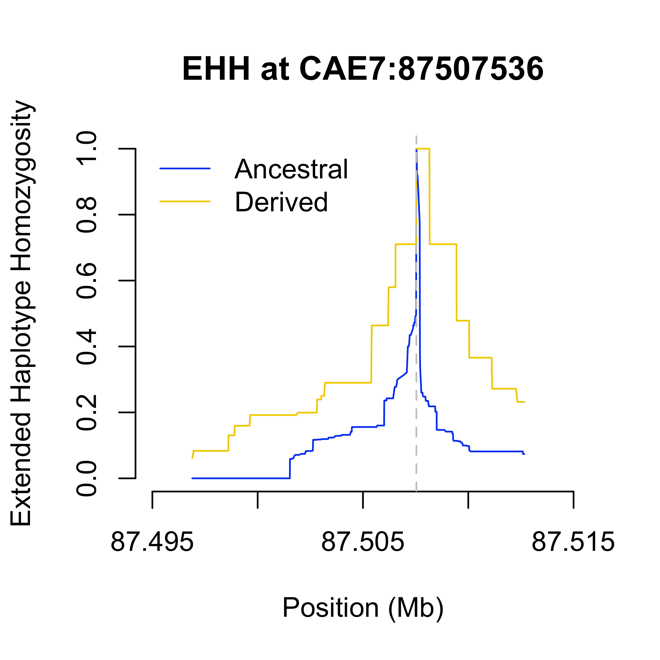

i) **87508112**

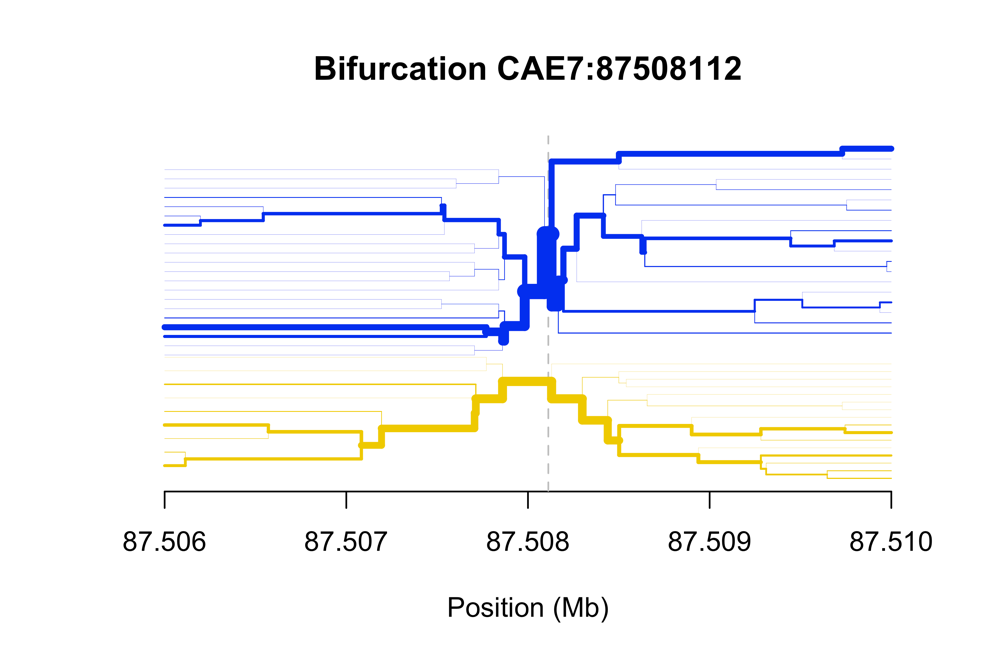

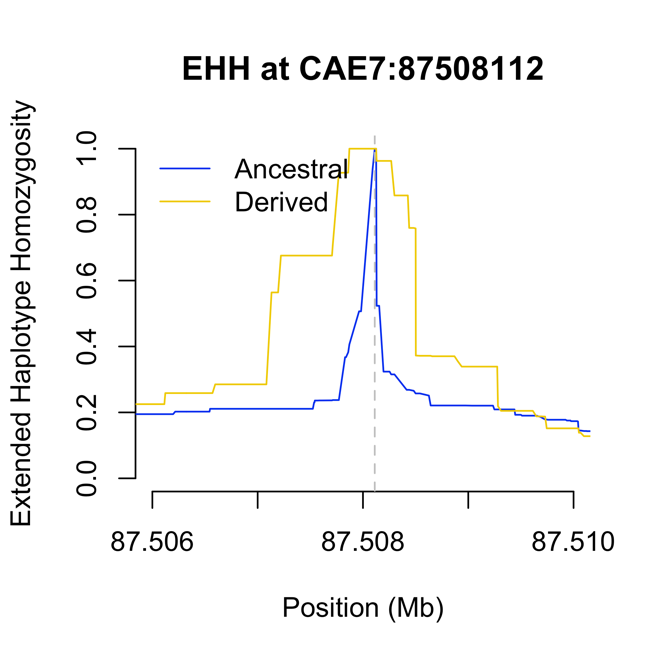

j) **87509248**

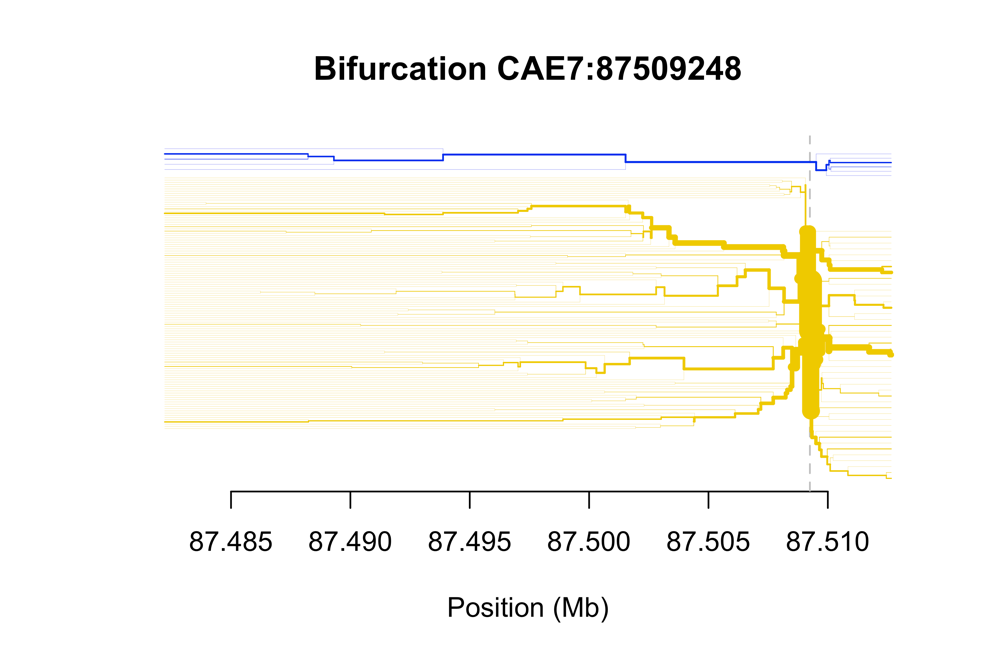

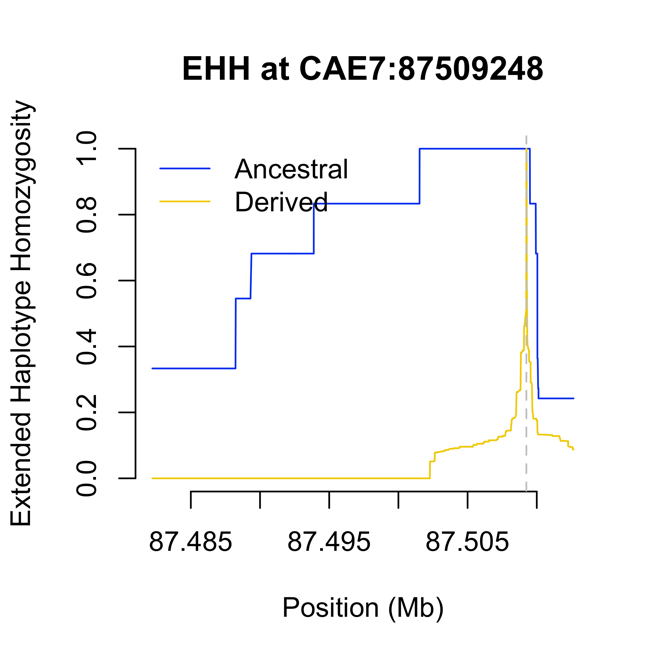

**Supplemental Figure 10**: OLS regressions showing relationship between derived allele frequency at loci experiencing putative selective sweeps and ecological variables significantly associated with derived allele frequency according to PGLS. Points in the figures use the same color convention as the rest of the paper.

a) **7:87491756** – note in these figures that the southern coastal corridor populations (KZN/EC) are clear outliers for both figures, and obviously driving the association. Were they removed, irradiance would not appear to be a factor in derived allele frequency for this SNP. Winter precipitation would show a clear pattern with the removal of EC/KZN.

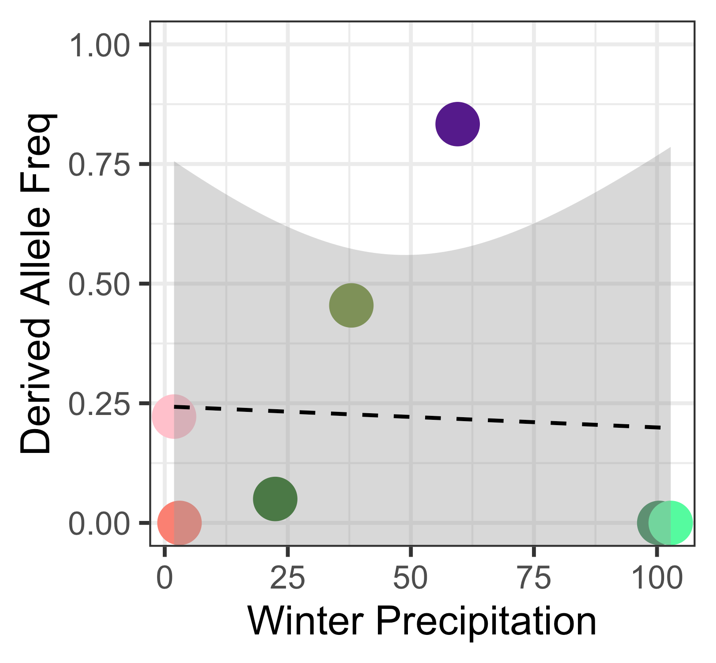

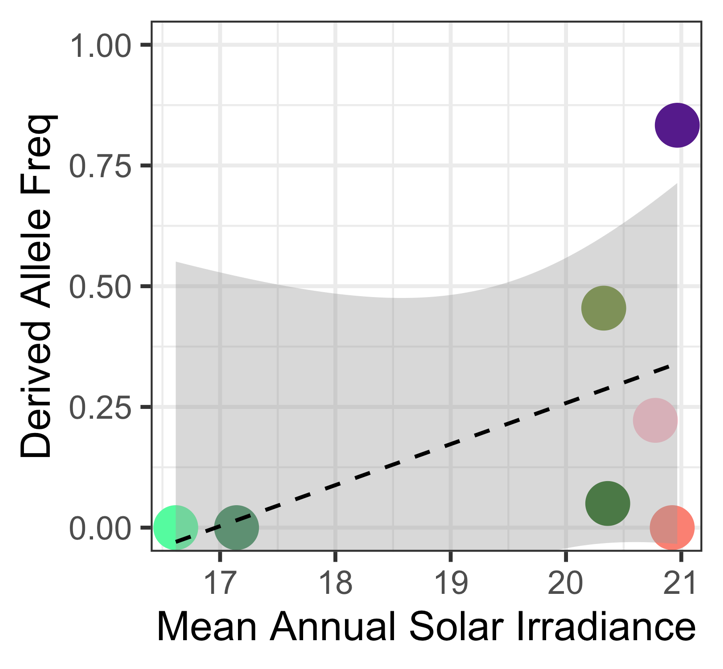

b) **7:87493023** – again the southern coastal corridor is driving the association with irradiance, although Free State populations also factor in. Notably, the EHH is around the derived allele at this locus, which is fixed in the non-*pygerythrus* populations.

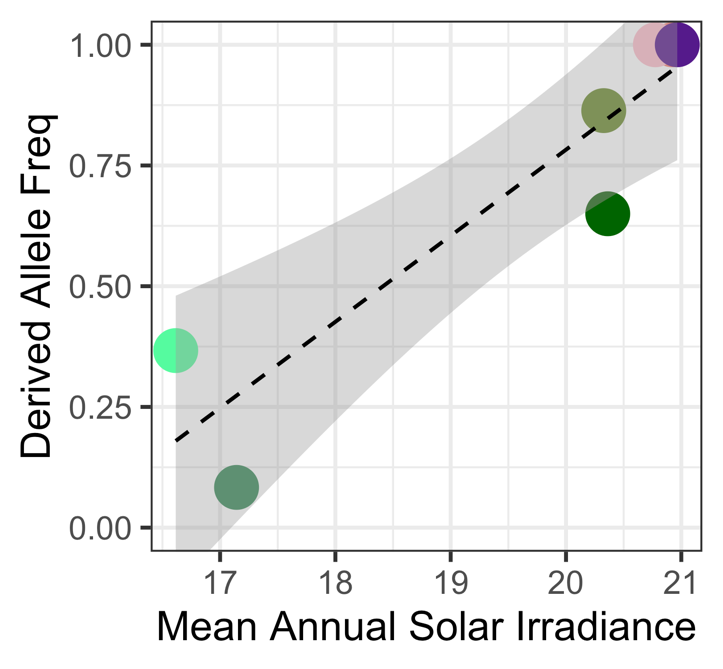

c) **7:87493613** – in this case, we have a relatively clear association of higher derived allele frequencies with lower mean winter precipitation. The southern coastal corridor represents an extreme of high precipitation in the winter, but not outliers.

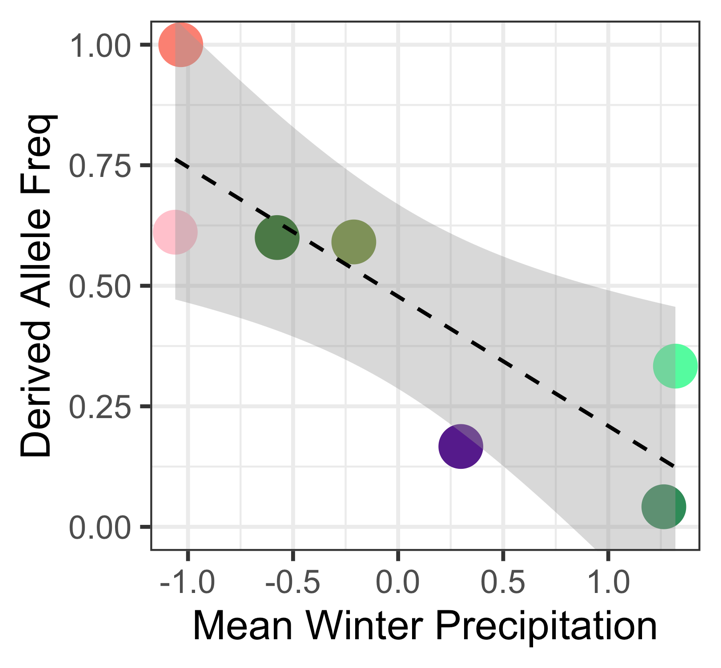

d) **7:87505287** – in this case, we again have a relatively clear association in which the southern coastal corridor represents an extreme but not outliers with respect to winter precipitation.

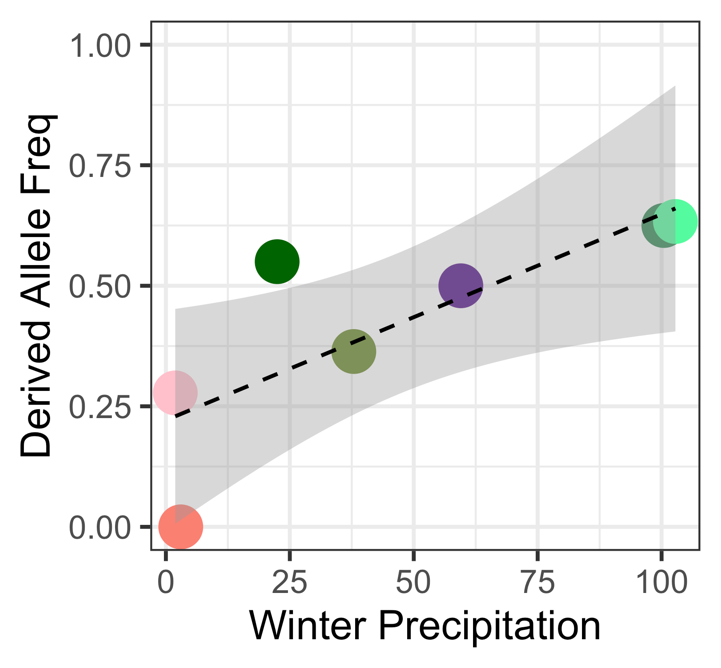

e) **7:87506414** – another locus for which the association with irradiance is driven by the southern coastal corridor populations.

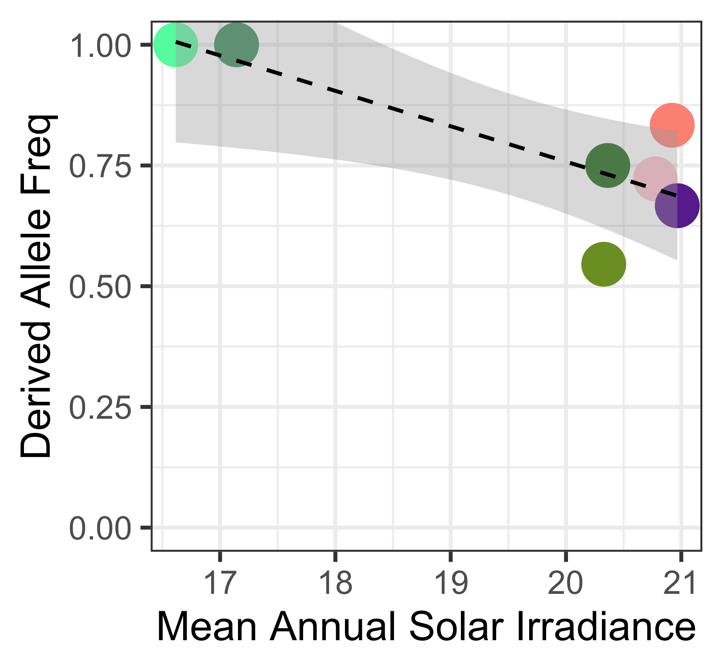

f) **7:87508112** – in this case, there’s a clear relationship with minimum temperature of the coldest month that is not influenced by the southern coastal corridor populations.

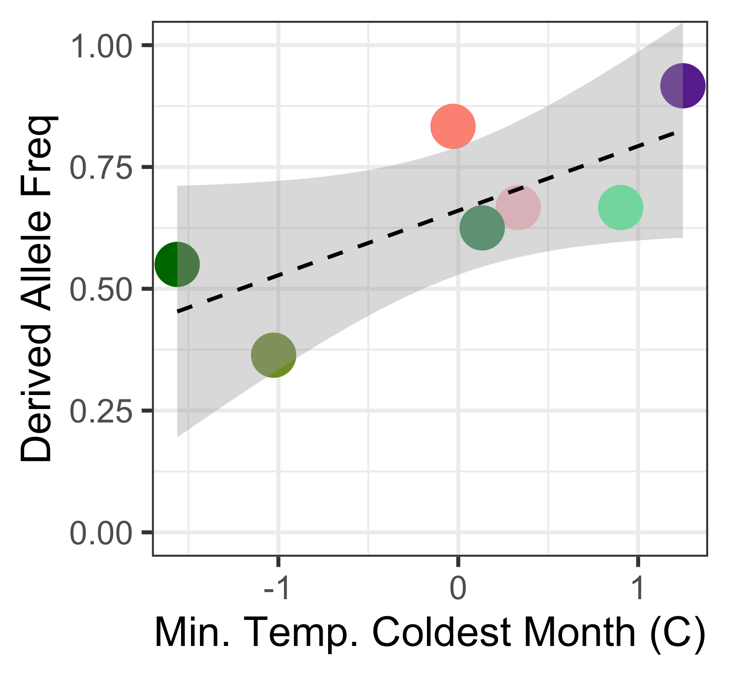

g) **7:87509248** – another case in which the southern coastal corridor populations are driving the association with irradiance.

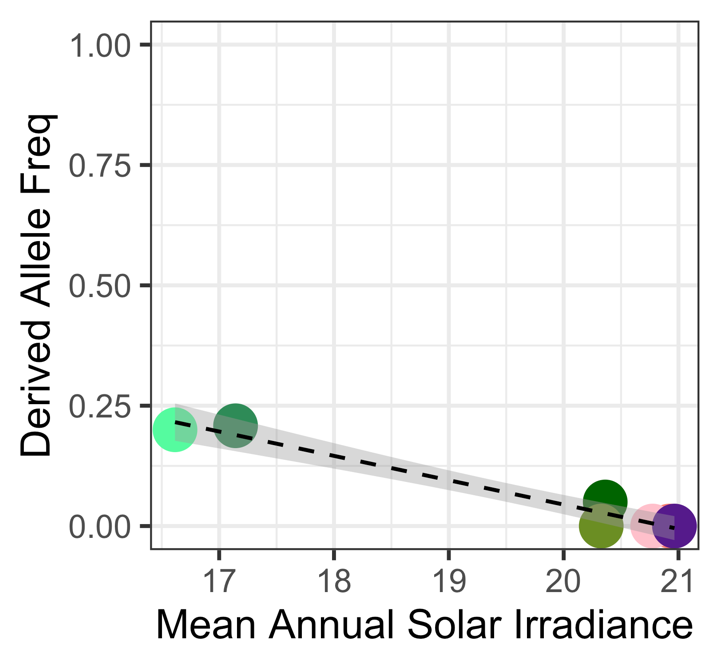
